# Neural markers of vulnerability to anxiety outcomes following traumatic brain injury

**DOI:** 10.1101/2020.04.20.051649

**Authors:** Juliana Popovitz, Shreesh P. Mysore, Hita Adwanikar

## Abstract

Anxiety outcomes following traumatic brain injury (TBI) are complex, and the underlying neural mechanisms are poorly understood. Here, we developed a multidimensional behavioral profiling approach to investigate anxiety-like outcomes in mice that takes into account individual variability. Departing from the tradition of comparing outcomes in TBI versus sham groups, we identified animals within the TBI group that are vulnerable to anxiety dysfunction by applying dimensionality reduction, clustering and post-hoc validation to behavioral data obtained from multiple assays for anxiety at several post-injury timepoints. These vulnerable animals expressed distinct molecular profiles in the corticolimbic network, with downregulation in GABA and glutamate, and upregulation in NPY markers. Indeed, among vulnerable animals, not resilient or sham controls, severity of anxiety outcomes correlated strongly with expression of molecular markers. Our results establish a foundational approach, with predictive power, for reliably identifying maladaptive anxiety outcomes following TBI and uncovering neural signatures of vulnerability to anxiety.

## INTRODUCTION

Anxiety-related disorders are a major outcome of traumatic brain injury (TBI) (Moore et al., 2006). They can develop months to years after the incident, with the pooled prevalence of anxiety estimated at 20% in the first year, and 36% at five years post-injury (Scholten et al., 2016). Notably, however, not all TBI patients are affected. Consequently, clinical studies aiming to address anxiety outcomes of TBI adopt rigorous exclusion criteria, such as structured interviews and scales, to determine which patients should be included in studies (Caspi et al., 2005, Al-Adawi et al., 2007).

By contrast, pre-clinical studies of TBI and anxiety-like behaviors have generally treated the injured population as a homogenous group, and compared the aggregate physiological and molecular sequelae in injured animals against those in control animals that experience sham surgery. These studies report a wide range of (conflicting) effects: from (i) a decrease in anxiety-like behaviors following a controlled cortical impact injury (Washington et al., 2012) and weight-drop injury (Pandey et al., 2009) at 20 to 30 days post-TBI, to (ii) an increase in anxiety-like behaviors following weight-drop (Meyer et al., 2012), lateral fluid percussion (Johnstone et al., 2015), and blast overpressure (Awwad et al., 2015), up to 3 months after later, as well as (iii) no change in anxiety-like behaviors in animals exposed to controlled cortical impact injury (CCI) (Sierra-Mercado et al., 2015), repeated weight-drop (Fidan et al., 2016) and lateral fluid percussion injury (Ferreira et al., 2014), up to 3 months after injury. Indeed, even when the injury model is fixed, the specific behavioral assay used to measure anxiety, the metric used to quantify it, as well as the time points of measurement, can all play a role in the results regarding anxiety outcomes following injury (Popovitz et al., 2019). This has made it difficult to identify reliably, the nature of anxiety dysfunction following TBI in pre-clinical models, as well as to probe the underlying neural mechanisms.

Inspired by the literature on the study of psychological stress (Cohen and Zohar, 2004, Cohen et al., 2006, Cohen et al., 2012), we considered whether the traditional approach in TBI studies of treating the injury group as a homogenous distribution may be obscuring the crucial factor of individual variability in behavioral outcomes of injury. Such variability has been observed in cognitive outcomes following TBI (Vonder Haar et al., 2016). In the stress literature, the role of individual differences in response to a common stressor has been defined via the idea of resilience (Smith et al., 2010, Wu et al., 2013), the organism’s ability to adapt to adversity. Exploiting this idea has led to insights about mechanisms underlying resilience and vulnerability to psychological stressors (Cohen et al., 2006, Cohen et al., 2012, Elliott et al., 2010).

Here, we set out to leverage the potential heterogeneity among individuals in anxiety responses to a CCI model of injury, in order to develop a pre-clinical approach analogous to the use of inclusion criteria in clinical TBI studies. Because of the inherent complexity of measuring anxiety in animal models, and the impact of different assays and timepoints (Popovitz et al., 2019), we reasoned that the approach in the stress literature of defining thresholds on pre-selected, unidimensional severity measures (‘cut-off behavioral criteria’ or CBC (Cohen and Zohar, 2004), would be ineffective for distinguishing vulnerability to anxiety responses following injury. In response, we developed an objective, multidimensional behavioral profiling, clustering and validation approach to identify vulnerable and resilient sub-groups of injured animals with distinct anxiety phenotypes. We then compared in vulnerable versus resilient groups, the expression of GABA (GAD 65/67), glutamate (VGLUT) and neuropeptide Y (NPY), key molecular markers (Heilig et al., 1989, Zhou et al., 2008, Lindell et al., 2010, Cohen et al., 2012), in three major components of the anxiety control network: the medial prefrontal cortex (mPFC), ventral hippocampus (vHPC), and basolateral amygdala (BLA). We found distinct signaling profiles for all three molecular markers in vulnerable animals, with broad reduction of GABA and VGLUT expression and broad increase in NPY expression, when compared to either resilient or sham animals. Notably, within the vulnerable group, molecular profiles correlated strongly with anxiety profiles. Moreover, ROC analysis revealed that a pre-injury anxiety measure was able to predict with ^~^70% accuracy, individuals that subsequently exhibit vulnerability to anxiety following injury. Our results demonstrate that this non-traditional, multidimensional profiling approach to analyzing dysfunction in anxiety-like behaviors within injured animals is powerful, and can reveal reliable neural markers of vulnerability. This work establishes a robust foundation for the dissection of neural mechanisms of anxiety outcomes following injury.

## RESULTS

### Multidimensional behavioral clustering with validation for identifying animals vulnerable to anxiety after TBI

Mice were tested on a battery of three commonly used behavioral assays for anxiety, namely, the elevated zero maze (EZM), open field test (OFT) and elevated plus maze (EPM) (Griebel et al., 1993, Belzung and Griebel, 2001, Prut and Belzung, 2003), before as well as after injury (Fig. 1A; Methods). The proportion of time that mice spend in the exposed zones of these behavioral arenas - open arms of the EZM and EPM, and the central zone of the OFT – is a standard metric used to quantify their anxiety-like behaviors (Griebel et al., 1993, Dawson and Tricklebank, 1995, Belzung and Griebel, 2001, Prut and Belzung, 2003, B. et al., 2017). We refer to it generally as ‘proportion of time spent in the anxiogenic zone’.

**Figure 1.**
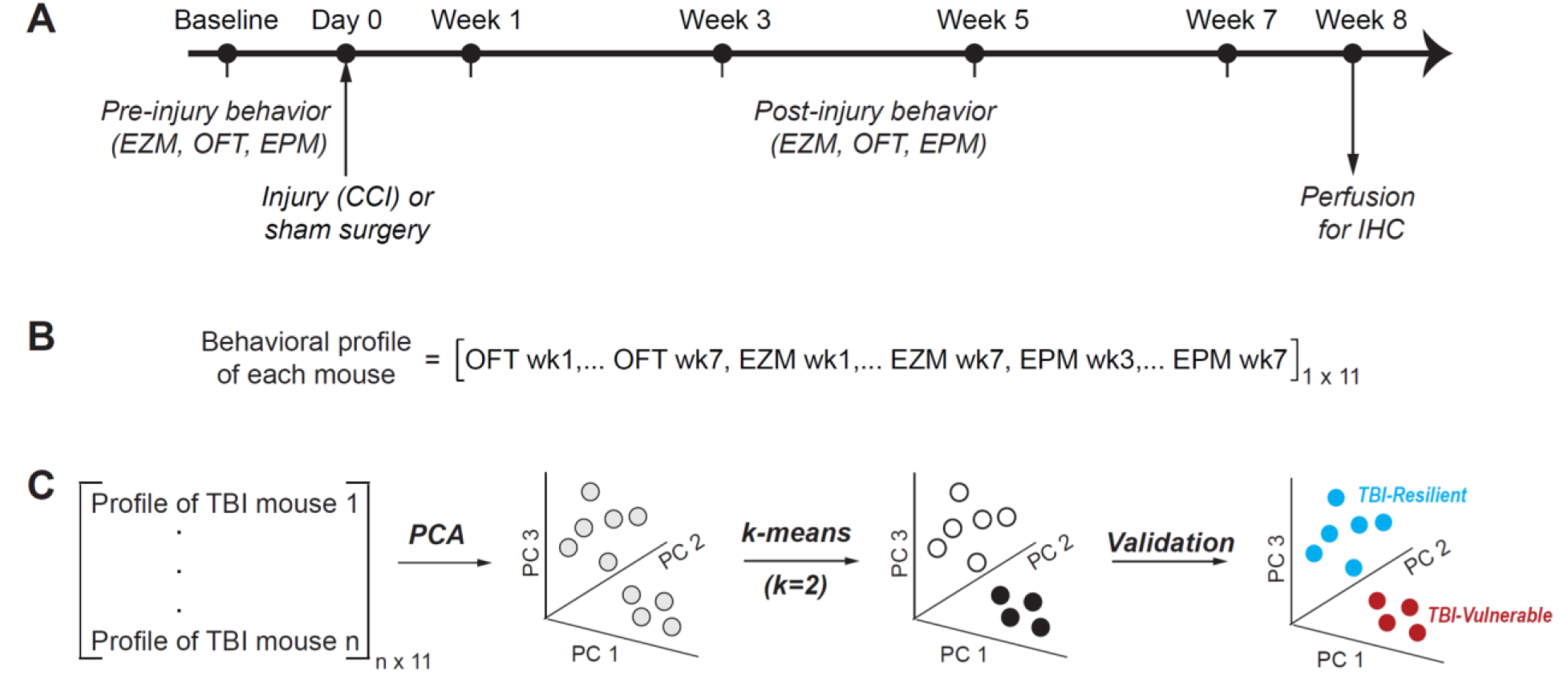
Multidimensional behavioral clustering with validation (MBCV) for identifying animals vulnerable to anxiety after TBI. **(A)** Experimental timeline and protocol for measuring anxiety-like behaviors before and after injury. EZM – elevated zero maze; OFT – open field test; EPM – elevated zero maze; all standard assays for measuring anxiety-like behaviors in rodents. CCI – Controlled cortical impact injury model. IHC – immunohistochemistry. **(B)** Anxiety behavioral ‘profile’ of a mouse: A 11-dimensional vector which combines behavioral data on the EZM, OFT and EPM across time points. Each element in the vector is the proportion of time spent by the mouse in the anxiogenic zones of one of the assays (ex. EZM) at one of the post-injury time points (ex. Week 7), normalized to the animal’s pre-injury baseline value in that assay (Methods). **(C)** Multidimensional behavioral clustering with validation approach: Behavioral profiles of mice exposed TBI are first subject to principal components analysis (PCA), and then to k-means clustering (with k=2) to identify two clusters in principal component (PC) space. The final validation step determines (i) whether animals in the two clusters exhibit distinct anxiety phenotypes after TBI, and if so, (ii) whether (and which) cluster consists of animals vulnerable to anxiety outcomes after TBI. This involves plotting and comparing between the clusters, anxiety metrics from each assay.

Metrics of anxiety-like behaviors as well as of general locomotion measured before injury were denoted as ‘baseline’ measurements; two pre-injury measurements were averaged to obtain the baseline values for each metric (Fig. 1A; Methods). A week following the baseline measurements, animals underwent either sham surgery (‘control’ group) or CCI injury (‘TBI’ group); mice were assigned randomly to these two groups. After another week of recovery, all animals were tested on the behavioral assays once every two weeks for a period of seven weeks following injury or sham surgery. Previous work has shown that such repeated testing in EZM does not produce a habituation effect, and whereas habituation is observed in OFT and EPM, it is consistent across sham and injury groups (Popovitz et al., 2019). For each animal, behavioral measurements were normalized to their respective baseline values to account for pre-surgery variability across animals. These (post-injury) measurements across mazes and time points yielded an 11-dimensional anxiety behavioral vector or ‘profile’ for each mouse (Fig. 1B; Methods). Mice were sacrificed at 8 weeks after injury, and immunohistochemical procedures performed on brain sections (Methods).

We asked if, based on their multidimensional anxiety behavioral profiles, mice that underwent TBI could be separated into two distinct groups such that one exhibited severe affective behavioral consequences of injury, and the other was largely resistant to effects of injury. To address this question, we developed a data-driven approach. We started with a cohort of 52 animals (Cohort A), of which 37 underwent TBI (and the rest underwent sham surgery; animals were assigned to conditions randomly; Methods). We performed principal components analysis (PCA) on the behavioral data from TBI animals (Fig. 1C; Methods), and found that 92% of the variance was explained by just 4 of the 11 dimensions (Fig. 1C; Fig. 2A-left, only 3 PCs shown). Next, we applied k-means clustering to the lower dimensional dataset of principal components to identify two distinct clusters (Fig. 1C; Fig. 2A-left, red and blue data; Methods). Since PCA transforms data points into a new coordinate frame, it was not possible to determine, directly, whether these clusters in PCA space represented low- and high-responders, and if so, which cluster represented what kind of response.

**Figure 2.**
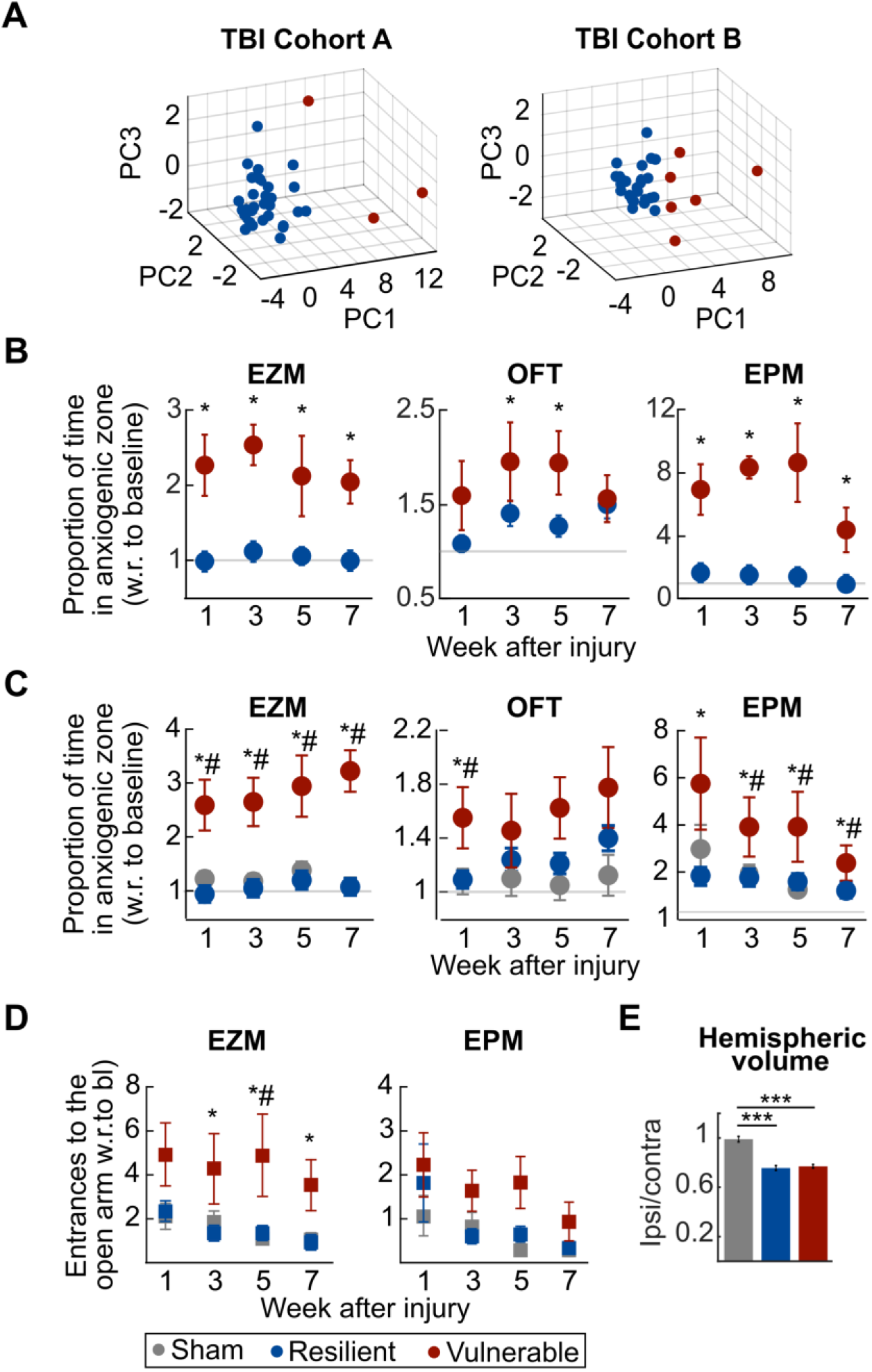
TBI animals present distinct behavioral profiles of vulnerability versus resilience to anxiety following TBI. **(A)** Multidimensional behavioral clustering applied to two different cohorts of mice that underwent TBI. Cohort A=37 TBI animals, Cohort B= 34 TBI animals; experiments on Cohorts A and B were performed ^~^1.5 years apart. Each dot is one mouse; shown are the first three PCs. The first four principal components captured most of the variability in both behavioral datasets - 91% for Cohort A and 87 % for Cohort B. Red and blue indicate the two clusters identified for each cohort. **(B**) Validation step (Cohort A). Plots of anxiety metric from each assay, comparing the red versus blue clusters of TBI animals from Cohort A (see text). <u>Left</u>: EZM; red animals shows decreased anxiety (greater proportion of time in anxiogenic zone) after injury compared to blue animals (2-way ANOVA: F=44.5, p=0, main effect.). <u>Middle</u>: OFT; 2-way ANOVA: F=9.47, p=0.002, main effect. <u>Right</u>: EPM; 2-way ANOVA: F=187.25, p=0, main effect. All three panels: Shown are mean +/- s.e.m; p<0.05, posthoc t-tests of red vs. blue by week, followed by Holm-Bonferroni multiple comparisons test (HBMC test). **(C)** Validation step applied to Cohorts A and B combined; red animals from both Cohorts were combined, and blue animals from both Cohorts were combined (see Supp. Fig. 1A and text). Gray data: Sham controls combined from both Cohorts (Cohort A: 15 sham controls, Cohort B: 17 sham controls). <u>Left</u>: EZM; red animals shows decreased anxiety after injury compared to blue and grey animals (3-way ANOVA: F=95, p=0, main effect.). <u>Middle</u>: OFT; 3-way ANOVA: F=8.37, p=0.0003, main effect. <u>Right</u>: EPM; 3-way ANOVA: F=12.45, p=0, main effect. All three panels: Shown are mean +/- s.e.m; and Ψ: p<0.05, posthoc t-tests of red vs. blue (or red vs. grey) by week followed by HBMC test. Results establish red animals as those vulnerable to anxiety outcomes following TBI, and red animals as resilient. **(D)** Total number of entrances into the open arms in the EZM and EPM; an independent metric of anxiety-like behavior that was not included in the multidimensional behavioral clustering with validation analysis (A-C); animals from Cohorts A and B combined. <u>Left</u>: EZM; red animals shows decreased anxiety after injury compared to blue and grey animals (3-way ANOVA: F=3.92, p=0.02, main effect.). ‘*’ and ‘#’: p<0.05, posthoc t-tests of red vs. blue (red vs. grey) by week followed by HBMC test. <u>Right</u>: EPM; 3-way ANOVA: F=1.68, p=0.18, main effect. Shown are mean +/- s.e.m. **(E)** Comparison of volume of injured brain hemisphere (normalized to contralateral hemisphere) between vulnerable, resilient and sham animals; estimated from post-hoc histology. (For sham control animals, ipsi and contra brain hemispheres were assigned randomly.) ANOVA: F=39.63, p=0.001 ‘***’: p<0.001, posthoc paired t-tests followed by HBMC test; resilient vs. vulnerable: p=0.67. Shown are mean +/- s.e.m. **See also Supp. Fig. 1.**

To address this issue and determine if animals in the two clusters exhibited distinct anxiety phenotypes after TBI, we plotted their anxiety metrics from each assay (Fig.2B). We found that animals in the red cluster showed a significant increase in the proportion of time spent in the anxiogenic zone when compared to animals from the blue cluster, and when compared to their baselines (Fig. 2B; repeated 2-way ANOVA, EZM: main effect: F=44.5, p=0; OFT: main effect: F=9.47 p=0.002; EPM: main effect: F=187.25, p=0). By contrast, animals in the blue cluster showed minimal change in anxiety metrics over time with respect to their pre-injury baseline (Fig. 2B; p>0.05; post-hoc t-tests followed by Holm-Bonferroni multiple comparisons test (HBMC test); Methods). Together, these results revealed that the two groups exhibited consistently distinct behavioral patterns across anxiety metrics and time following injury. Moreover, they demonstrated that animals in the blue cluster were largely ‘resilient’ to TBI, with behavior not different from their baselines, whereas animals in the red cluster were ‘vulnerable’ to anxiety dysfunction following injury, with behavior substantially different from their baseline, as well as from that of resilient animals. In this cohort, we found that just 3/37 animals were vulnerable to anxiety dysfunction following injury. We refer to this analysis method of identifying distinct behavioral groups from multidimensional profiles as ‘multidimensional behavioral clustering with validation’ (MBCV).

To increase the size of the vulnerable sample for subsequent analyses, we repeated the experiment on a different cohort of 51 animals (Cohort B), of which 34 underwent TBI (and the rest underwent sham surgery). We applied our MBCV analysis approach to the behavioral data from TBI animals in this cohort (Fig. 2A; right panel; first four principal components account for 87% variability; Methods). We identified 6 vulnerable and 28 resilient mice in this cohort; their behavioral data are shown in Supp. Figure 1A. For all subsequent analyses, we merged the vulnerable (as well as resilient) groups from the two cohorts.

First, we compared the proportion of time spent by all animals vulnerable to TBI, all animals resilient to TBI, as well as all sham animals in the anxiogenic zone in each of the behavioral assays (Fig. 2C). We confirmed systematic differences between the vulnerable and both resilient and sham animals, but no differences between the resilient and sham animals (Fig. 2C; EZM: repeated three-way ANOVA, main effect of treatment, F=95, p=0, post-hoc: vulnerable versus resilient and vulnerable versus sham, p<0.01 in all time-points. OFT: repeated three-way ANOVA, main effect of treatment, F=8.37, p=0.0003, post-hoc: vulnerable versus resilient and vulnerable versus sham p<0.05 on week one. EPM: repeated three-way ANOVA: main effect of treatment, F=12.45, p=0, effect of time, F=4.05, p=0.007, post-hoc, vulnerable versus resilient, p<0.05 in all time-points, vulnerable versus sham, p<0.05 on weeks three, five and seven). Specifically, vulnerable animals spent a significantly higher proportion of time exploring the open arms in both the EPM and the EZM, and a trend towards greater exploration of the center in the OFT, as compared to the resilient animals (Fig. 2C). By contrast, in the EZM and EPM, resilient and sham animals showed no changes from baseline in the proportion of time spent in the anxiogenic zone (p>0.05, post-hoc t-test followed by HBMC test; Methods).

To further validate our approach, we investigated whether vulnerable and resilient animals exhibited similar anxiety outcomes when assessed on a different anxiety metric, one that was not used in the MBCV analysis to identify these groups. To this end, we analyzed the number of entrances to the open arm for the EZM and the EPM (Fig. 2D), and found similar results to those from proportion of time spent in the anxiogenic zone. In the EZM, vulnerable animals exhibited a greater number of entrances into the open arm than resilient and sham animals (Fig. 2D, left: repeated three-way ANOVA, F=3.92, p=0.02), with significant differences observed between vulnerable and resilient animals on weeks three, five and seven (p<0.05, post-hoc t-tests followed by HBMC test; Methods), and significant differences between vulnerable and sham animals on weeks five and seven (p<0.05, post-hoc t-tests followed by HBMC test; Methods). In the EPM, there is no significant difference between groups (repeated three-way ANOVA, F=1.68, p=0.18), but there was a trend indicating that vulnerable animals exhibited more entrances into the open arm compared to resilient and control animals. Thus, findings based on our MBCV approach were robust to the anxiety metric used.

We assessed potential deficits in general locomotion by measuring the total distance travelled by mice in each of the three assays (Supp. Fig 1B). There was no difference in locomotion (normalized to baseline) between the groups in the EZM (repeated three-way ANOVA, F=0.17, p=0.85,). In the EPM, there was a main effect of injury (repeated three-way ANOVA, F(2)=3.84, p=0.02,), but no significant post-hoc effect. In the OFT, there was a main effect of injury (repeated three-way ANOVA, F=29.74, p=0,) with vulnerable (but not resilient) animals being significantly more active than sham controls on all weeks (p<0.05, post-hoc t-tests followed by HBMC test; Methods). These results are in agreement with previous findings of increased overall locomotor activity following injury in the open field test (Budinich et al., 2013, B. et al., 2017, Corrigan et al., 2017). Notably, however, the lack of an effect on locomotion in the EZM and EPM, along with enhancement, rather than impairment, in locomotion in the OF, suggests that it is not a confounding factor for the observed differences in anxiety-like behavior between the vulnerable and resilient groups.

We next wondered whether differences in injury severity between groups might account for their anxiety phenotypes. To this end, for each animal, we assessed the volume of the injured hemisphere (ipsi) relative to the volume of the uninjured hemisphere (contra) and compared across groups (Fig. 2E; Methods). We found a reduction in the normalized (ipsi/contra) hemispheric volume in the peri-injury area in both the TBI groups (vulnerable and resilient), compared to sham controls (three-way ANOVA, F=39.63, p<0.001, effect of treatment; p<0.001, post-hoc t-test followed by HBMC test; Methods, vulnerable and resilient vs. sham). However, there was no difference between vulnerable and resilient animals (t-test, p=0.67), ruling out differences in injury as a source of vulnerability. Additionally, there was no effect on the contralateral peri-injury hemispheric volume in TBI animals compared to sham controls (Supplementary Table 1, three-way ANOVA, F=1.52, p=0.23).

Our approach to identify distinct sub-groups within TBI animals was motivated by the hypothesis that the distributions of behavioral measures following injury versus sham surgery may overlap significantly, thereby making it difficult to isolate reliable differences in behavioral outcomes (and neural mechanisms) by the standard approach of comparing between these groups. To test the veracity of this underlying hypothesis, we compared the distributions of the proportion of time spent in the open arm of the EZM between the TBI and sham groups. We found that, indeed, the distributions of anxiety behavioral metrics in the TBI and sham control groups exhibited substantial overlap (Supp. Fig. 1C). Taken together, the above results establish the MBCV as a robust, data-driven approach for identifying two groups of animals, vulnerable and resilient, with distinct anxiety-like behavioral outcomes following injury.

### Clustering sham animals using the MBCV approach does not produce behaviorally distinct groups

The success of MBCV approach applied to TBI animals raised the question as to whether this approach is effective specifically for TBI animals, or whether the act of splitting any distribution (i.e., any batch of animals) into two groups can yield groups with distinct anxiety profiles, simply by virtue of the intrinsic variability within the distribution. We tested this question as follows. Since each cohort also included animals that underwent sham surgery and for which the same behavioral measures were collected as for TBI animals, we applied the MBCV approach to data from sham animals. To this end, we applied PCA and 2-means clustering to each sham cohort independently (Fig 3A), just as we had done for TBI animals (Fig. 2A). We then examined the individual anxiety metrics for the two clusters from each cohort, and found no systematic differences between them (Supp. Fig. 2). For improved statistical power of this comparison, we also combined clusters across cohorts, in both possible ways (Fig. 3A: dark green + teal and orange + maroon, or dark green + maroon and orange + teal). We found no systematic differences between the resulting groups, in either case (Fig. 3B, dark green + teal vs. orange + maroon: EZM, F=0.95 p=0.33; EPM, F=1.14 p=0.28; Fig. 3C, dark green + maroon and orange + teal: EZM, p=0.33, F(2)=0.95; OFT, p=0.003, F=8.65, brown animals show higher values than pink animals; EPM, p=0.007, F=7.37, brown animals show lower values than pink animals, no post hoc effect). In other words, MBCV applied to sham animals did not produce groups that exhibited distinct patterns of anxiety outcomes that were consistent across anxiety metrics. These results demonstrated that injury caused dysfunctional anxiety profiles in vulnerable animals, and that the MBCV method is able to detect them.

**Figure 3.**
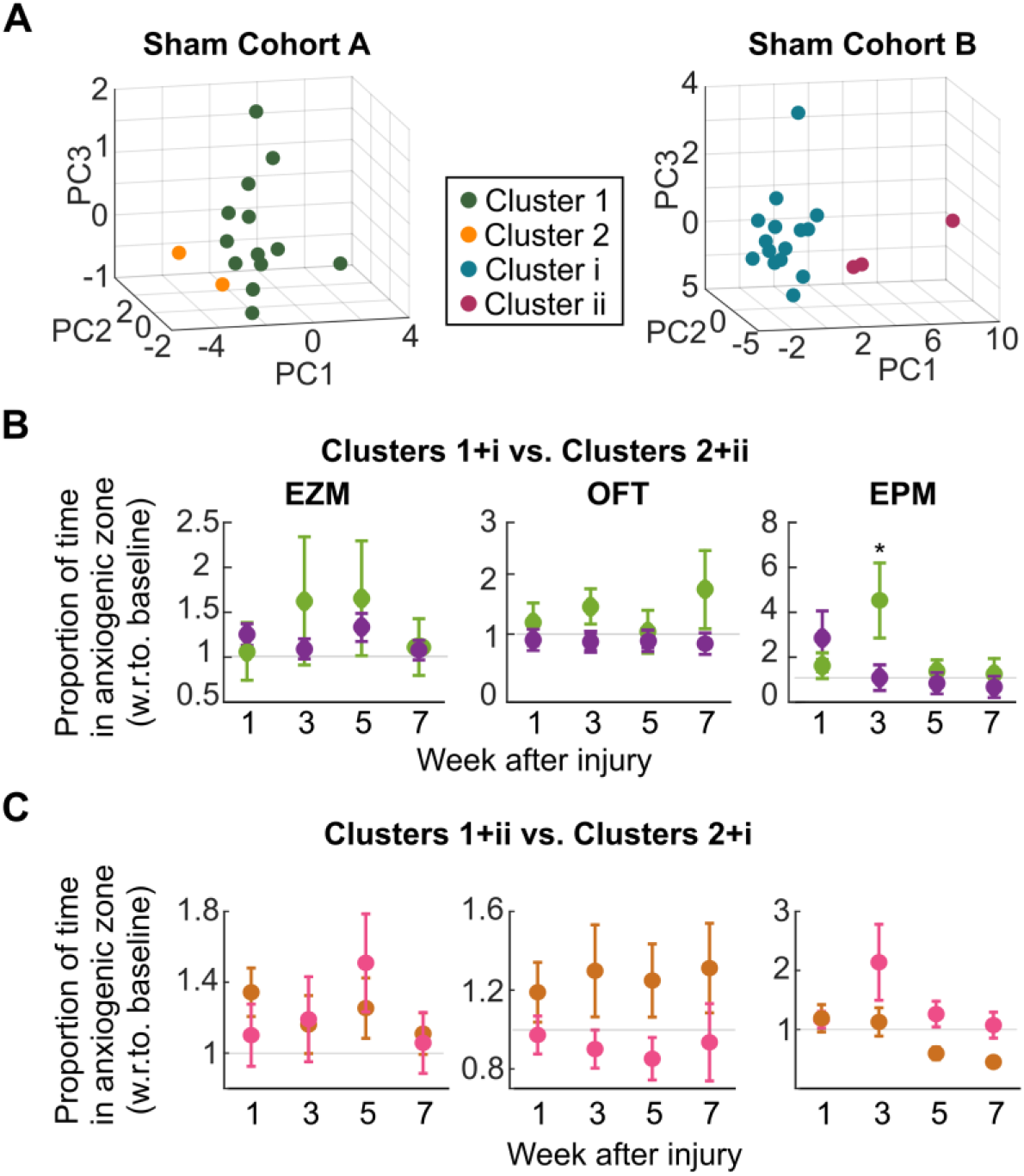
Sham animals do not present distinct behavioral profiles of vulnerability versus resilience to anxiety dysfunction. **(A)** Multidimensional behavioral clustering (as in Figure 2A) applied to animals that underwent sham surgery in each of the Cohorts (Cohort A=15 sham animals, Cohort B= 17 sham animals; within each cohort, animals assigned to sham versus TBI groups randomly). Colors indicate the two clusters identified among sham animals in each cohort; all other conventions as in Figure 2A. **(B,C)** Validation step applied to sham clusters from Cohorts A and B combined in both possible ways: (B) Dark green cluster (Cohort A) and teal cluster (Cohort B) combined versus orange cluster (Cohort A) and maroon cluster (Cohort B) combined. <u>Left:</u> EZM; no main effect of group (2-way ANOVA, F=0.95, p=0.33). <u>Middle:</u> OFT; no main effect of group (2-way ANOVA, F=1.14, p=0.28). <u>Right:</u> EPM; 2-way ANOVA, F=8.8 p=0.003; with post-hoc effect on week 3 (p<0.05). (C) Dark green cluster (Cohort A) and maroon cluster (Cohort B) combined versus orange cluster (Cohort A) and teal cluster (Cohort B) combined. <u>Left:</u> EZM; no main effect of group (2-way ANOVA, F=0, p=0.98). <u>Middle:</u> OFT; main effect of group (2-way ANOVA, F=8.65, p=0.003); but no post-hoc effects (p>0.05). <u>Right:</u> EPM; main effect of group (2-way ANOVA, F=7.37, p=0.007), direction of effect in opposite from that in OFT (middle); no post-hoc effects (p>0.05). Results demonstrate that multidimensional behavioral clustering applied to sham animals does not produce groups with consistently distinct anxiety profiles. **See also Supp. Fig. 2.**

### Vulnerable animals present downregulation of GAD and vGLUT, and upregulation of NPY markers in the mPFC

Having identified vulnerable versus resilient groups based on their anxiety behavioral profiles, we next asked if there were systematic molecular differences that could serve as neural signatures of vulnerability and resilience. To assess molecular differences, we perfused the brains of the animals and performed immunohistochemistry against three markers - an inhibitory marker (GAD65/67), an excitatory marker (VGLUT), and neuropeptide Y (NPY). We did this in the mPFC, a key brain region involved in the control of anxiety (Adhikari, 2014). Figure 4A shows the location of imaging and sample images for three markers across animal groups (resilient, vulnerable and sham).

**Figure 4.**
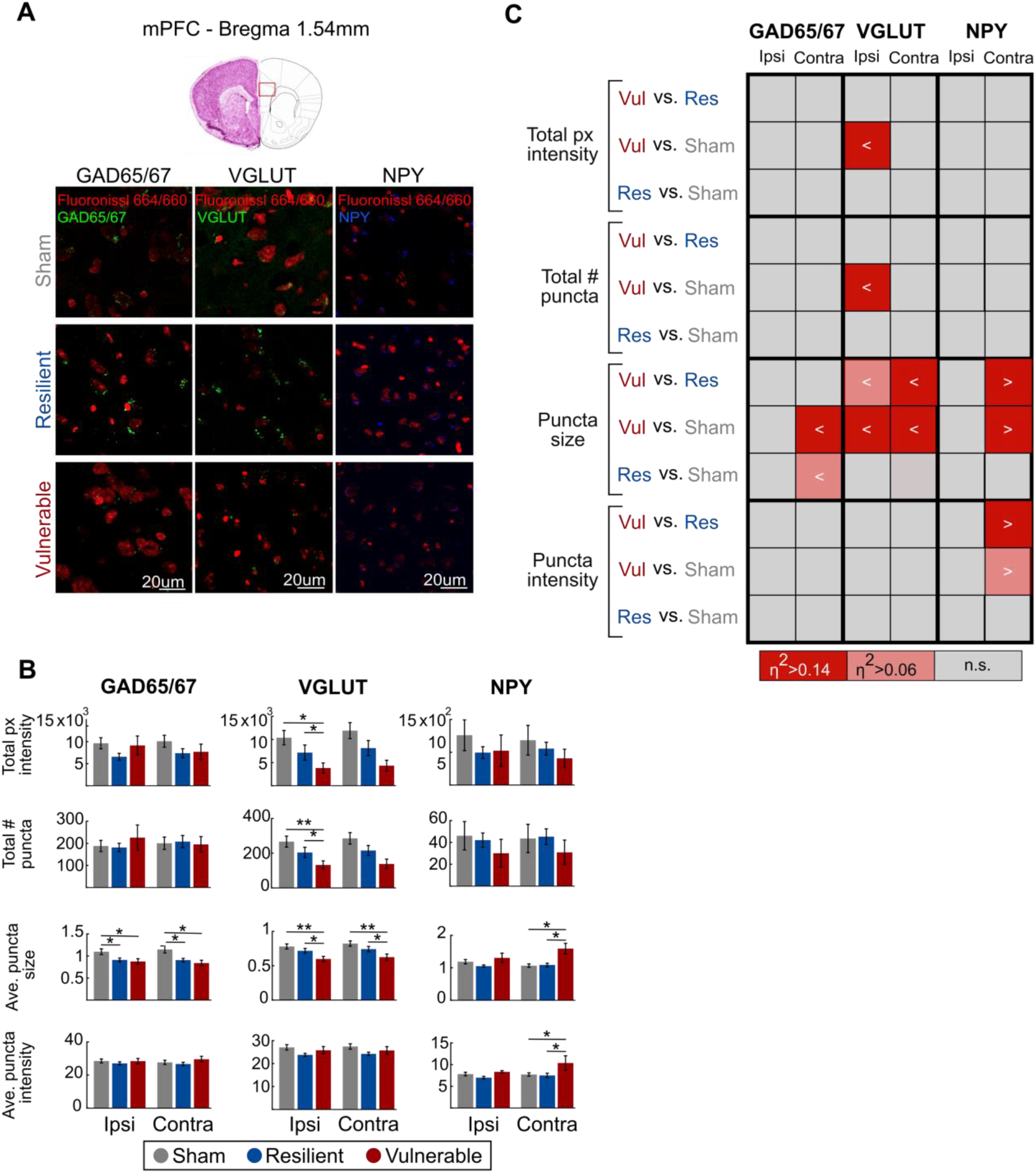
Vulnerable animals present downregulation of GAD and vGLUT, and upregulation of NPY immunostaining in the mPFC. **(A)** <u>Top</u>: Schematic of brain section showing where mPFC images were taken. <u>Bottom</u>: Example sections from sham, resilient and vulnerable animals (rows) showing immunostaining for GAD65/67, vGLUT and NPY (columns). **(B)** Comparison of various metrics of immunostaining (rows) for each of the three molecular markers (columns) in mPFC. Shown are mean +/- s.e.m; number of animals: vulnerable: all 9, resilient: GAD65/67: n=28, VGLUT: n=29, NPY: n=23; sham: GAD65/67: n=16, vGLUT: n=17, NPY: n=12; resilient and sham animals for immunostaining were selected randomly from their respective groups) *: p<0.05 and **: p<0.01, for difference between groups: ANOVA followed by post-hoc t-tests with HBMC (see also text). **(C)** Visualization of effect sizes (η^2^) for comparisons between groups (vulnerable, resilient, sham) of the different metrics of immunostaining in mPFC that presented a statistically significant difference (post-hoc: p<0.05) (from B). Colors indicate size of effect. For each comparison of x vs. y, < (>) indicates that x<y (x>y).

Staining for GAD in the mPFC showed a downregulation in both resilient and vulnerable animals compared to sham controls. There was a significant decrease in the average size of GAD 65/67 puncta in the ipsilateral as well as contralateral sides (Fig 4B- left column, third row; ANOVA, F=4.07, p=0.02; contra: ANOVA, F=6.54, p=0.003; post-hoc t-test followed by HBMC test; Methods, p<0.05 for both sides of the brain). However, there were no differences in GAD staining between vulnerable and resilient animals.

By contrast, staining for VGLUT in the mPFC showed a downregulation, specifically in vulnerable animals, when compared to resilient animals as well as sham controls. The total intensity of VGLUT pixels was marginally lower, and number of their puncta and the average puncta size were significantly lower on the ipsilateral side in vulnerable animals (Fig. 4B-middle column, top three rows; pixel intensity: ANOVA, F=2.31, p=0.1; puncta number: ANOVA, F=3.35, p=0.04, post-hoc t-test followed by HBMC test; Methods p<0.05; puncta size: ANOVA: F=3.44, p=0.04, post-hoc t-test followed by HBMC test; Methods, p<0.05). The average puncta size was lower on the contralateral side as well for vulnerable animals in comparison to resilient animals and sham controls (Fig. 4B - middle column, third row; ANOVA: F=3.4, p=0.04, post-hoc t-test followed by HBMC test; Methods, p<0.05). Notably, in all cases, there were no differences between the resilient animals and sham controls.

Staining for NPY, a marker for resilience to stress (Lindell et al., 2010, Cohen et al., 2012), also showed a change that was specific to vulnerable animals, but interestingly, showed an upregulation in the mPFC. The average puncta size and intensity were significantly higher on the contralateral side in vulnerable animals (Fig. 4B - right column, bottom two rows; size: ANOVA, F=8.99, p=0.0007; intensity: F=3.53, p=0.03). Notably, in both cases, vulnerable animals were different from both control and resilient animals (post-hoc t-test followed by HBMC test; Methods, p<0.05). There were no differences between resilient and sham animals.

To visualize these effects succinctly, we plotted a matrix of effect sizes observed in the comparisons between all pairs of groups (vulnerable vs resilient, vulnerable vs sham, and resilient vs sham) for each of these molecular measures (Fig. 4C; Methods); the size of the effect was coded by color. This matrix indicates clearly that the resilient group does not show strong changes with respect to control (η^2^>0.06) in any of the metrics measured. By contrast, the vulnerable group shows a strong effect size (η^2^>0.06) with respect to sham control in metrics such as puncta size of GAD 65/67 staining, puncta size, number and total pixel intensity of VGLUT immunostaining, and puncta size of NPY immunostaining (GAD: vulnerable vs sham, puncta size, contra: η^2^=0.22, p=0.01; resilient vs sham, puncta size, contra: η^2^=0.22, p=0.01; VGLUT: vulnerable vs sham, puncta size, ipsi: η^2^=0.3, p=0.003, contra η^2^=0.29, p=0.004; number of puncta, ipsi: η^2^=0.27, p=0.006; total px intensity, ipsi: η^2^=0.24, p=0.01; NPY: vulnerable vs sham, puncta size, contra: η^2^=0.37, p=0.008). Interestingly, when comparing vulnerable and resilient groups, there is a moderate or strong effect on VGLUT immunostaining (VGLUT: vulnerable vs resilient, puncta size, ipsi: η^2^=0.33, p=0.001, contra: η^2^=0.16, p=0.02), as well as NPY immunostaining (NPY: vulnerable vs resilient, puncta size, contra: η^2^=0.33, p=0.001; puncta intensity contra: η^2^=0.14, p=0.04).

### Vulnerable animals present significant differences in molecular markers compared to resilient and sham animals in the BLA and vHPC

Additionally, in these same animals, we also measured expression of GAD65/67, VGLUT and NPY in two other nuclei important for the control of anxiety-like behaviors, namely the basolateral amygdala (BLA) and ventral hippocampus (vHPC) (Adhikari et al., 2010, Felix-Ortiz et al., 2013). Figure 5A and C show the location of imaging in each of these brain regions.

**Figure 5.**
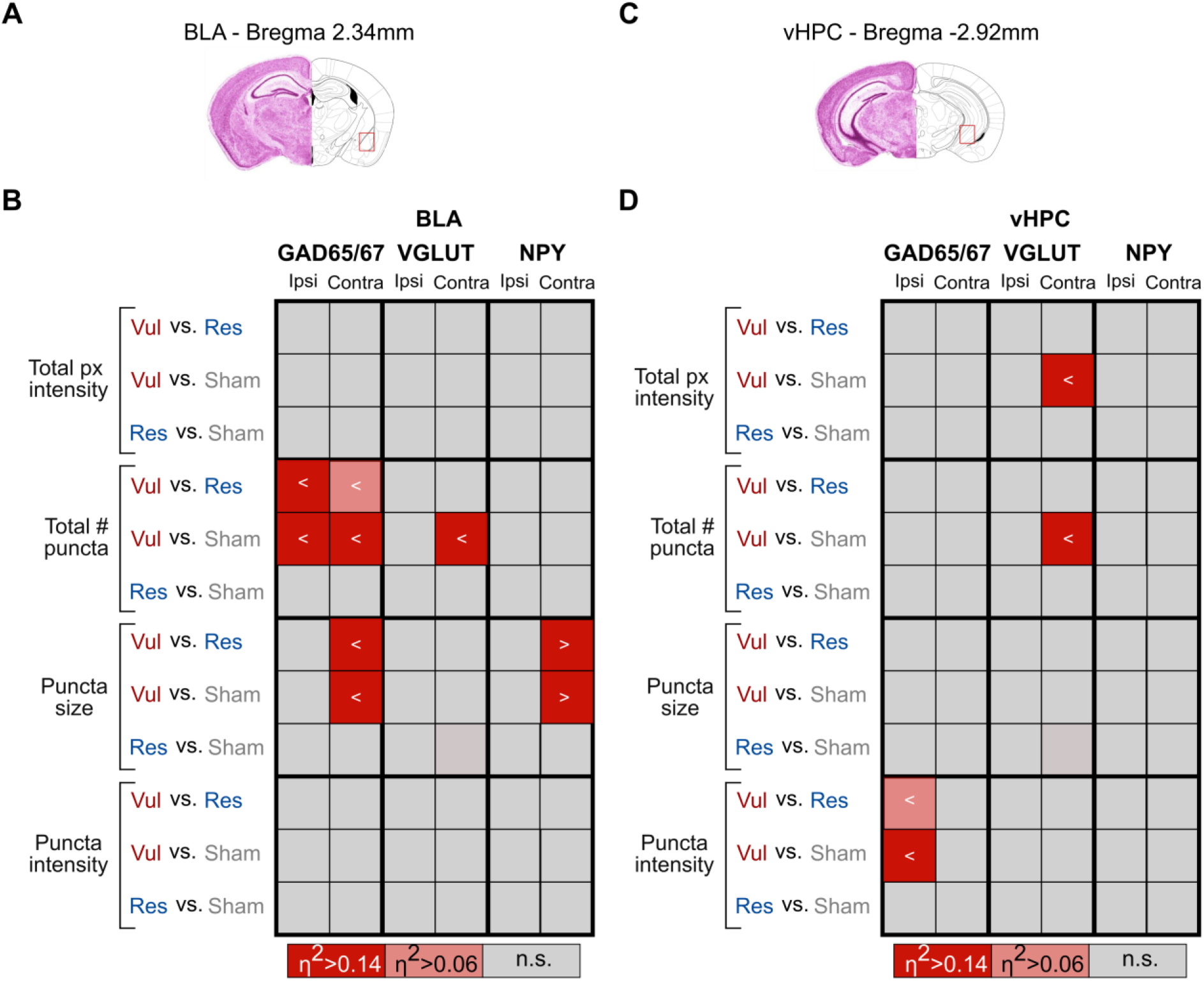
Vulnerable animals present significant differences in molecular markers compared to resilient and sham animals in the BLA and vHPC. **(A,C)** Schematic of brain sections showing where BLA (A) and vHPC (C) images were taken. **(B,D)** Visualization of effect sizes (η^2^) for comparisons between groups (vulnerable, resilient, sham) of the different metrics of immunostaining in BLA (B) and vHPC (D) that presented a statistically significant difference (post-hoc: p<0.05) (from Supp. Fig. 3). Colors indicate size of effect. For each comparison of x vs. y, ‘<‘ indicates that x<y. Number of animals: vulnerable: all 9, resilient: GAD65/67: n=28, VGLUT: n=29, NPY: n=23; sham: GAD65/67: n=16, vGLUT: n=17, NPY: n=12. **See also Supp. Fig. 3.**

In the BLA, fewer IHC metrics than in the mPFC showed significant effects of animal group (vulnerable, resilient or sham), but the overall nature of effects was similar (Supp. Fig. 3A – plots of IHC metrics and Fig. 5B - plot of effect sizes). Specifically, vulnerable animals, when compared to resilient (and sham) animals showed a moderate reduction in the total number of GAD 65/67 puncta in the ipsilateral side (Supp. Fig. 3A, GAD column, 2^nd^ row; ANOVA: F=3.91, p=0.02; post-hoc t-test followed by HBMC test; Methods, p<0.05; Fig. 5B), and a moderate decrease in the size of GAD puncta in the contralateral side (Supp. Fig. 3A, GAD column, 3^rd^ row; ANOVA: F=4.37, p=0.01, post-hoc t-test followed by HBMC test; Methods, p<0.05; Fig. 5B). For VGLUT staining, vulnerable animals showed a significant reduction in the total # VGLUT puncta compared to sham controls on the contralateral side (Supp. Fig. 3A, VGLUT column, 1^st^ row; ANOVA: F=2.24, p=0.02, post-hoc t-test followed by HBMC test; Methods, p<0.05; Fig. 5B), indicating a downregulation of VGLUT. However, we found no significant differences specifically between vulnerable and resilient animals on any of the VGLUT metrics (Supp. Fig. 3A, and Fig. 5B, VGLUT column; Fig. 5B). For NPY staining, vulnerable animals showed a strong increase in the size of NPY puncta compared to resilient (as well as control) animals on the contralateral side (Supp. Fig. 3A, NPY column, 3rd row; ANOVA: F=8.78, p=0.001, post-hoc t-test followed by HBMC test; Methods, p<0.05; Fig. 5B). Notably, there were no observed differences in the BLA between resilient animals and sham controls in any of the IHC metrics (Fig. 5B).

In the vHPC as well, only a few IHC metrics showed an effect of the animal group (Supp. Fig. 3B – plots of IHC metrics and Fig. 5D – plot of effect sizes), but the effects were broadly consistent with those in the other areas. Specifically, we found that vulnerable animals, when compared to resilient animals, showed a significant reduction in the intensity of GAD puncta (ipsilateral side; Supp. Fig. 3B, GAD column, 4^th^ row; ANOVA: F=3.88 p=0.03, post-hoc t-test followed by HBMC test; Methods, p<0.05; Fig. 5D). Additionally, vulnerable animals, when compared to sham controls, showed a reduction in GAD staining (Supp. Fig. 3B, GAD column, 4^th^ row, ipsilateral puncta intensity: post-hoc t-test followed by HBMC test; Methods, p<0.05; Fig. 5D), and a reduction in VGLUT staining (Supp. Fig. 3B, VGLUT column, 1^st^ row, contralateral total intensity: ANOVA: F=3.13, p=0.05; 2^nd^ row, contralateral total # puncta: ANOVA: F=3.36, p=0.04, post-hoc t-test followed by HBMC test; Methods, p<0.05; Fig. 5D). Notably, there were no observed differences in the vHPC between resilient animals and sham controls in any of the IHC metrics (Fig. 5D).

Thus, when comparing the vulnerable versus resilient groups, in the BLA, there was a moderate downregulation of GAD staining and a strong upregulation of NPY staining, but no effects of VGLUT staining. In the vHPC, there was a weak downregulation of GAD, and no effects on VGLUT or NPY expression.

Together with the results from mPFC, our findings reveal that animals that are vulnerable to anxiety outcomes following injury exhibit distinct molecular profiles as compared to resilient injured animals (or sham controls), with stronger overall effects in the mPFC than vHPC or BLA: decreases in metrics of VGLUT and GAD 65/67 staining in mPFC, decreases in metrics of GAD65/67 staining in BLA and vHPC, as well as increases in NPY staining in mPFC and BLA. We refer to these molecular metrics (total of 17) that show a selective difference in vulnerable versus resilient and control animals, as molecular signatures of the vulnerability to TBI dependent anxiety dysfunction.

### Molecular metrics in vulnerable animals correlate with behavioral anxiety outcomes

The finding of distinct molecular signatures for vulnerable individuals motivated us to ask, whether molecular profiles varied systematically with the extent of vulnerability to anxiety outcomes. To examine this question, we tested for correlation between molecular metrics that showed significant effects in vulnerable animals, and a behavioral measure of anxiety. We reasoned that behavioral measurements made on week 7, the last week of testing before perfusion and immunohistochemistry, would constitute the most relevant ones for testing the correlation. Additionally, of the different week 7 measurements, we reasoned that the proportion of time spent in the open arms of the EZM, the metric that exhibited the largest and most consistent effect of vulnerability (Fig. 2C), represented the most plausible candidate.

Examination of the relationship between this behavioral measure of anxiety and each of the 17 molecular indicators of vulnerability post-injury revealed that nearly half of the indicators (8/17) showed a significant correlation within the vulnerable animals (Fig. 6A. Specifically, we found that there was a strong correlation between VGLUT immunostaining in the mPFC (total intensity, puncta number and size), and in the BLA (total intensity), with individual vulnerability (Fig. 6A). Similarly, there was a strong or moderate correlation between GAD 65/67 observed in the mPFC (puncta size), in the BLA (puncta size and number), and in the vHPC (puncta intensity), with individual vulnerability (Fig. 6A). These correlation effects are summarized as a color-coded matrix in Figure 6B.

**Figure 6.**
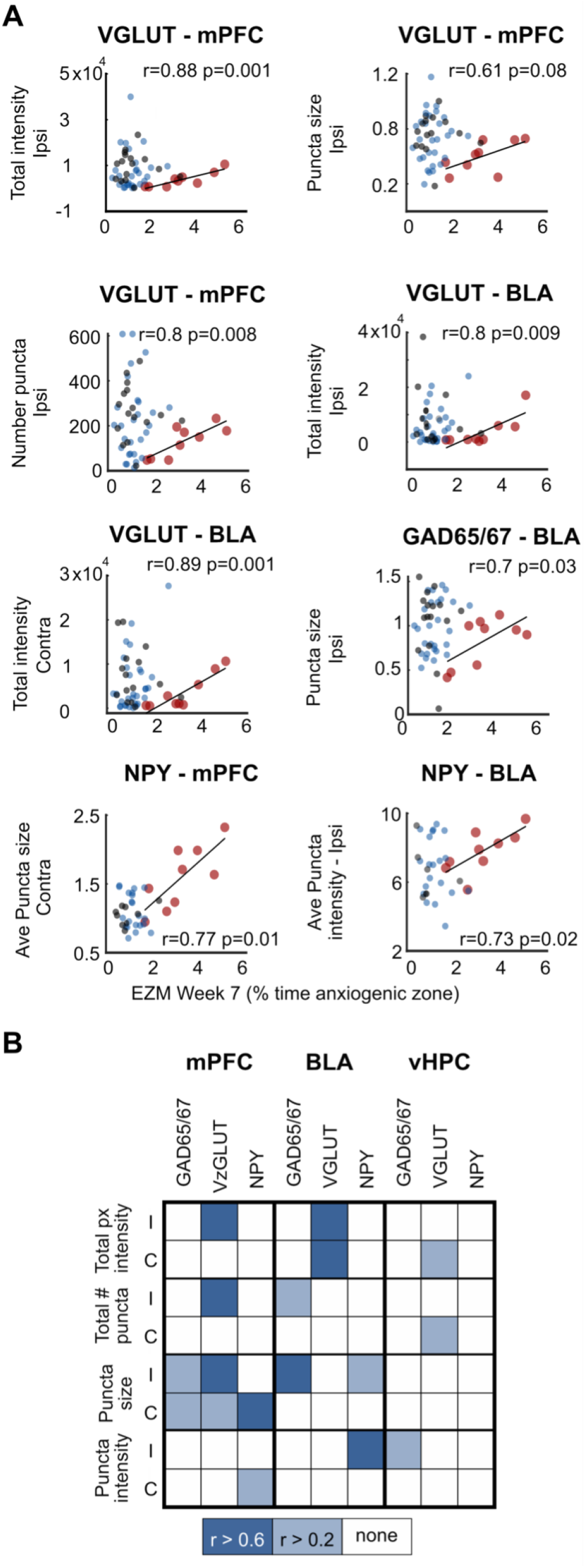
Molecular metrics in vulnerable animals correlate with anxiety outcome measured in week 7 (EZM) **(A)** Scatter plots of molecular metrics versus anxiety metric (proportion of time spent in the anxiogenic zone) measured on week 7 in the EZM. Each dot denotes one mouse; red- vulnerable animals (all 9); blue - resilient animals (GAD65/67: n=28, vGLUT: n=29, NPY: n=23), grey - sham controls (GAD65/67: n=16, vGLUT: n=17, NPY: n=12). r: Pearson’s correlation, line: best-fit line, p: p-value from Pearson’s correlation test. Plots are shown for the 8 molecular metrics that exhibited a significant correlation (for vulnerable animals) with the anxiety metric; all 17 molecular metrics that showed a significant difference between vulnerable and resilient or sham animals (from Figures 4 and 5) were tested. **(B)** Visualization of the strength of correlations in A. The matrix represents strength of correlations between the proportion of time spent by vulnerable animals in the anxiogenic zone of EZM in week 7, against each of various molecular metrics (rows) of GAD65/67, VGLUT and NPY (columns) measured in the three brain regions in the ipsi (I) and contra (C) hemispheres. All correlation values indicated are positive. **See also Supp. Fig. 4.**

To test if this strong correlation was unique to vulnerable animals, or whether the variability intrinsic to measurements made within any group of animals could show such a correlation, we repeated the correlation analysis above for resilient animals as well as sham controls (Fig. 6A, blue and grey dots, respectively). We found that there were no significant correlations between the molecular indicators and the behavioral metric for animals in either of these two groups, highlighting the significance of this relationship in vulnerable animals.

The above results largely held true when we used a different behavioral metric to examine correlations with molecular indicators, although fewer metrics exhibited strong correlations (Supp. Fig. 4). These results indicated that the existence of correlation between molecular markers and behavioral measures of vulnerability is general, and indicate EZM wk7 as the most informative single behavioral indicator of vulnerability.

### Pre-injury measurements can predict post-injury vulnerability

Lastly, we wondered whether the pre-injury behavior of the vulnerable individuals could predict their vulnerability to anxiety dysfunction post injury. To test this, we compared the baseline measurements of anxiety- like metrics for animals from these groups. We found that the vulnerable group did not show differences from either the resilient or sham groups for either the EPM or the OFT (three-way ANOVA; OFT: F=1.01, p=0.36; EPM: F=1.81, p=0.16,). However, in the EZM, vulnerable animals spent less time in the open arm during baseline, compared to the resilient animals (Fig 7A-left, EZM: F=4.97, p=0.008).

**Figure 7.**
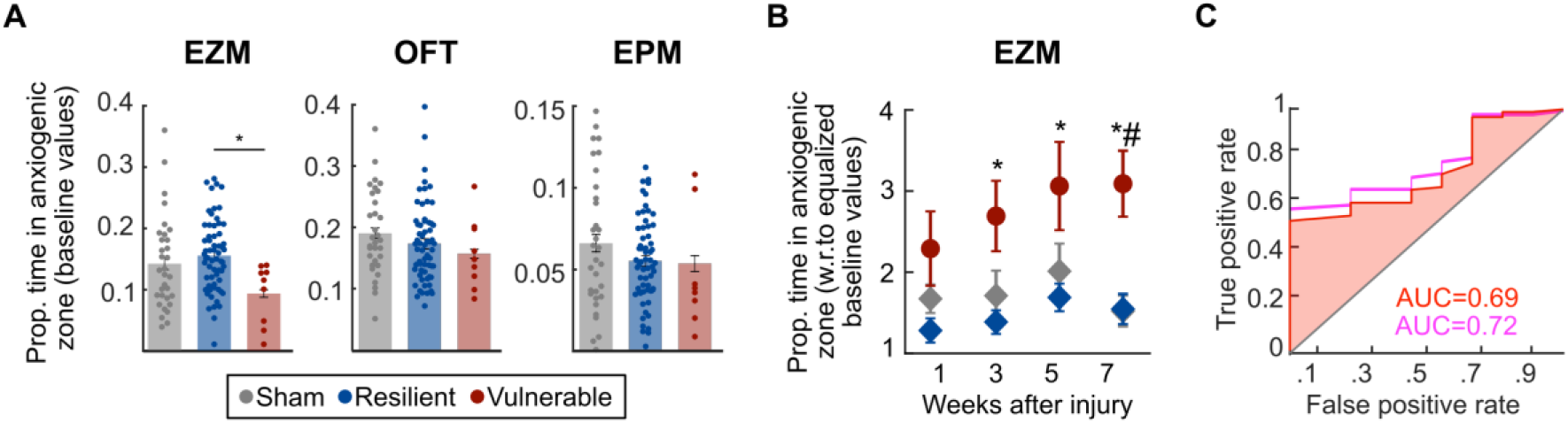
Predicting vulnerability to post-injury anxiety dysfunction using pre-injury measurements. **(A)** Anxiety metrics measured with different assays at baseline (pre-injury). Left: EZM; main effect of group (ANOVA; F=4,97, p=0.008), with vulnerable animals showing reduced proportion of time (increased anxiety) compared to resilient and sham animals (p<0.05, post-hoc t-tests with HBMC). Middle: OFT; no effect of group (ANOVA; F=1.01, p=0.36). Right: EPM; no effect of group (ANOVA; F=1.81, p=0.16). **(B)** Plots of anxiety metrics from EZM post-injury, computed after artificially setting baseline means of resilient and sham groups to be equal to the baseline mean of the vulnerable group (‘baseline equalization’; Methods). Diamonds: data after normalization with respect to equalized baselines. Vulnerable animals continue to show anxiety dysfunction (increased exploration of the anxiogenic zone) following injury compared to resilient and sham control animals (3-way ANOVA: F=60.03, p=0.002, main effect). (Ψ): p<0.05, posthoc testing by week of red vs. blue and red vs. grey with HBMC. Shown are mean +/- s.e.m. **(C)** ROC analysis across all animals (TBI and sham; red; Methods) or only on TBI animals (pink) on anxiety metric measured with EZM at baseline (pre-injury). Area under ROC curves (AUC) = 0.69 (red); 0.72 (pink); results demonstrate the efficacy of pre-injury values in predicting post-injury vulnerability to anxiety.

Since the anxiety dysfunction in vulnerable animals presents as an increase in the proportion of time spent in the anxiogenic zone after injury (when normalized to baseline), we wondered whether, in the case of EZM, the observed decrease in baseline values presents a confound: could the normalization by the smaller baseline values for vulnerable animals be the driving factor for the observed increase in their post-injury metrics? To test this decrease, we artificially set the average baseline values for resilient and sham groups to match that of the vulnerable animals (by subtracting the difference in mean baseline values, Methods). Following this baseline ‘equalization’, we compared the post-injury outcomes normalized to these adjusted baseline values among the three groups. We found that vulnerable group continued to show a significant increase in time spent in the anxiogenic zone compared to the resilient or sham groups (Fig. 7B), ruling out differences in baseline values as a confound for the observed post-injury effects.

These results, together with the fact that pre-injury baseline values for vulnerable animals were no different from those of resilient animals in other assays (OFT and EPM), showed that the enhanced exploratory behavior in the vulnerable group, compared to the resilient and sham groups, is largely, if not entirely, the consequence of injury.

Having established that the pre-injury decrease in the EZM metric was not a confound, we wanted to explore further the implications of this decrease. We asked if the pre-injury anxiety measure was predictive of vulnerability post-injury. To test this, we performed an ROC analysis (Methods) on the preinjury EZM measurements and found that vulnerable animals could be identified with 69% accuracy among all animals (Fig. 7C, red), and with 72% accuracy among all injured animals (Fig. 7C, pink), based solely on baseline measurements of anxiety-like behaviors. Thus, pre-injury differences in EZM behavior are predictive of vulnerability to anxiety dysfunction following TBI, and represent a starting point for understanding pre-existing neural mechanisms underlying this vulnerability.

## DISCUSSION

Anxiety outcomes following traumatic brain injury are complex and variable. The prevalence of anxiety disorders is higher among TBI patients than the general population (Rao and Lyketsos, 2002). At the same time, following TBI, not all patients are equally affected in terms of neuropsychiatric outcomes, with anxiety disorders developing in about a third (Scholten et al., 2016). In this study, we hypothesized that TBI similarly leads to a range of anxiety outcomes in an animal model of injury, and that these outcomes correlate with different neuro-molecular signatures. To test this hypothesis, we developed a robust multidimensional behavioral clustering and validation (MBCV) approach that could be applied to anxiety metrics from several assays (EZM, OFT and EPM) and at various time points (up to 7 weeks) following injury in mice (focal CCI). This approach demonstrated that animals can be reliably clustered into two groups based on their multidimensional behavioral profiles: Vulnerable animals showed significantly increased exploration of the anxiogenic zone following injury, whereas resilient animals displayed no change in their anxiety-like behavior and were similar to sham-injured controls. Additionally, it revealed that in regions of the brain associated with anxiety processing(Adhikari et al., 2010, Tye et al., 2011), not only was vulnerability was accompanied by the distinct downregulation of VGLUT and GAD65/67, and upregulation of neuropeptide Y, but also that these molecular metrics exhibited strong correlations with individual behavioral vulnerability. Finally, we also found that postinjury vulnerability to anxiety could be predicted with ~70% accuracy by a single pre-injury behavioral metric of anxiety, namely, by the proportion of time spent in the anxiogenic zone of the EZM in baseline. Our work establishes MBCV as a powerful approach for quantifying anxiety dysfunctions of TBI and elucidating the underlying neural mechanisms reliably in mouse models.

### Behavioral identification of a vulnerable profile

The concept of psychological vulnerability or resilience has been extensively described in humans, in the context of resilience to psychosocial stress (Rutter, 1993, Cicchetti, 2010, Luthar et al., 2014) and in animal models of stress (Cohen and Zohar, 2004, Cohen et al., 2007, Cohen et al., 2008). Resilience is defined broadly as the ability of an organism to adapt to adversity, by the engagement of coping strategies (Smith et al., 2010). Vulnerability, on the other hand, is the lack of coping strategies and has been implicated in triggering or worsening several psychiatric disorders, such as schizophrenia (Nuechterlein and Dawson, 1984), post-traumatic stress disorder (Resnick et al., 1992), depression (Abramson et al., 2002) and anxiety (Williams et al., 2005). The role of resilience and vulnerability in the context of TBI has received little explicit attention, and few studies have measured patients resilience following injury (White et al., 2010, McCauley et al., 2013, Bushnik et al., 2015, Holland and Schmidt, 2015, Kreutzer et al., 2016). Even fewer studies have applied the concept of resilience and vulnerability to outcomes of TBI in pre-clinical models. In one study, cognitive deficits following bilateral frontal CCI were examined (Vonder Haar et al., 2016), and three (distinct) groups of animals were identified (resilient, vulnerable, and chronically-impaired). However, this grouping was based on using preselected cut-off criteria applied to unidimensional metrics. To the best of our knowledge, no previous study has applied the concept of resilience and vulnerability to (multidimensional) anxiety outcomes of TBI in animal models.

Our primary hypothesis was that affective behavioral outcomes can show vulnerability or resilience to injury, leading to increased variability in the observed behavior (Washington et al., 2012, Almeida-Suhett et al., 2014, Tucker et al., 2017). We found substantial overlap in the distributions of anxiety outcomes between TBI versus sham injured animals, supporting this hypothesis, and also demonstrating that standard approach of comparing outcomes between TBI and sham groups may be flawed. Moreover, a multidimensional behavior profile has been found to be important in characterizing affective behavioral responses following injury (Popovitz et al., 2019), which makes the application of simple criteria-based thresholds to separate behavioral subsets difficult. (Cohen et al., 2004, Elliott et al., 2010). In this study, we addressed these challenges by developing a quantitative approach that can be applied to high dimensional behavioral datasets (assays x time-points) to identify distinct behavioral subgroups within the TBI animals. It detects distinct behavioral subsets based on an objective data-driven method, rather than by using pre-determined criteria. Using this approach, we identified resilient and vulnerable subsets within the TBI injured group. Notably, application of the same approach to sham-surgery animals did not result in groups that displayed consistently distinct anxiety outcomes. This indicates that the MBCV approach is sensitive to the injury-induced emergence of distinct behavior profiles; it does not identify two distinct anxiety groups in any arbitrary population of animals. Importantly, this suggests that the emergence of post-injury behavioral dysfunction is driven by injury-induced activation of specific neural mechanisms. Identifying these mechanisms can allow us to begin addressing these behavioral outcomes of injury in the vulnerable population.

Our approach necessarily involved repeated testing: this permitted the inclusion of the time-course of behavioral deficits in the analysis. A general concern with repeated behavioral testing is habituation to the testing arena, leading to potential confounds in the interpretation of behavioral changes/outcomes: changes could be associated with shifts from unconditioned to learned fear response (Bertoglio and Carobrez, 2000). However, in mice following TBI, previous work has shown that for repeated testing at two-week intervals, habituation effects do not present a confound (Popovitz et al., 2019): In EZM and OFT, no perceptible habituation effects were observed, and in EPM, habituation effects have been shown to impact both TBI and sham animals equally. Our results, here, show similar effects, indicating that habituation cannot account for the relative increase in exploration of the anxiogenic zone by the vulnerable group. Instead, results in this study suggest that the diversity in the behavioral response following injury is crucial factor in understanding the behavioral and mechanistic consequences of injury.

### Increased exploration of exposed spaces: adaptive or dysfunctional?

Vulnerable animals present an increased exploration of the exposed (high anxiety) zones within various behavioral arenas following TBI, which can be interpreted as a decrease in anxiety-like behavior. Consistent with the fact that avoidance of potential threat is a fundamental aspect of human anxiety (Borkovec et al., 2004), increased exploration of the exposed zone, i.e., a reduction in anxiety, is often interpreted as an adaptive response, while decreased exploration, i.e., an increase in anxiety, is interpreted as a maladaptive response (‘anxiety disorder’). However, engaging in an excessive exploration of a dangerous area can expose animals to unnecessary threats (Machado et al., 2009, Moscarello and Maren, 2018), demonstrating lack of behavioral control, and therefore, can itself be maladaptive. This could also indicate a deficit in risk-assessment, or an increase in risk-taking behavior. Clinical studies have demonstrated increased impulsivity, compulsive behaviors, addiction, careless or violent behavior and suicide, in patients with moderate to severe injury (Greve et al., 2001, Grados et al., 2008, Noggle and Pierson, 2010, Fecteau et al., 2013, James et al., 2014, Juengst et al., 2014), consistent with deficits in risk-assessment. We conclude that post-injury increase in exploration of the exposed zones by vulnerable animals represents a maladaptive anxiety state.

### Molecular signatures of individual vulnerability to anxiety

Identification of behaviorally distinct subsets within the injury group allowed us to then probe distinctions in the underlying molecular signaling within key brain areas in the cortico-limbic circuit associated with anxiety-like behaviors: the mPFC, BLA, and vHPC (McIntosh et al., 1987, McIntosh et al., 1989, Pierce et al., 1998, Sanders et al., 1999, Creeley et al., 2004, Thompson et al., 2005, Adwanikar et al., 2011, Reneer et al., 2011, Wang et al., 2011, Si et al., 2013, Yang et al., 2013, Cernak, 2015). Specifically, we focused on molecular markers related to the excitatory-inhibitory (E-I) balance.

In the mPFC, activation of GABA receptors (Shah et al., 2004) as well as activation of pyramidal neurons are both known to increase anxiety-like behaviors (Berg et al., 2019). Similarly, in the BLA, overall hyperexcitability (Prager et al., 2016) as well as increased firing rates of inhibitory neurons (Lee et al., 2017, Babaev et al., 2018) are associated with enhanced anxiety-like behaviors. Additionally, activation of vHPC neurons projecting to the mPFC has been shown to increase anxiety-like behaviors (Parfitt et al., 2017). Taken together, an increase in both excitation as well as inhibition in key hubs for anxiety control are associated with increases in anxiety. These results have been framed in the context of the excitatory-inhibitory (E-I) balance, with changes in E-I balance leading to dysfunctional anxiety behaviors.

Following TBI, however, the signaling changes in the corticolimbic circuit associated with anxiety outcomes are poorly understood. In the mPFC, whereas E-I signaling changes linked to cognitive and fear dysfunction following injury have been explored (Kobori and Dash, 2006, Schneider et al., 2016), those associated with anxiety behaviors are less clear. In the BLA, results regarding GABAergic and glutamatergic signaling are contradictory, with some studies reporting increases and other reporting decreases following injury (Ajao et al., 2012, Reger et al., 2012, Malkesman et al., 2013, Almeida-Suhett et al., 2014, Palmer et al., 2016, Popovitz et al., 2019, Beitchman et al., 2020). In the hippocampus, dysfunction of inhibitory synapses and reduction in glutamate precursors have been reported following injury (Witgen et al., 2005, Cole et al., 2010). Together with the conflicting anxiety-behavioral outcomes reported after injury, the association of molecular signaling mechanisms with changes in anxiety-related behavior following injury remain unclear.

Our results, here, demonstrate that animals vulnerable to anxiety dysfunction following injury, compared to resilient animals, exhibit down-regulation of VGLUT and GAD signaling in all three brain regions - mPFC, BLA and vHPC. Considering that vulnerable animals show decreased anxiety-like behaviors, our findings provide a clear link, for the first time, between dysfunctional anxiety following injury and the underlying molecular changes, one that is consistent with the well-established molecular mechanisms for the regulation of anxiety behaviors.

Resilience/vulnerability to behavioral dysfunction is a complex, multifaceted phenomenon, dependent on the interaction of both genetic and environmental factors (Reichmann and Holzer, 2016). A molecular marker that has been associated with resilience is Neuropeptide Y (NPY). It has been identified as a marker of resilience in outcomes to stress exposure (Flierl et al., 2009, Koliatsos et al., 2011, Rubovitch et al., 2011). Specifically, in the context of affective dysfunction, exogenous NPY or NPY overexpression in the hippocampus or the amygdala, is associated with anxiolytic behavior or adaptive mechanisms (Thorsell et al., 1999, Primeaux et al., 2005, Sajdyk et al., 2008).Consistent with these reports, we find increased expression of NPY in mPFC and the BLA, specifically in the vulnerable group, which exhibit decreased anxiety-like behaviors. In this context, injury can be considered a ‘stressor’, exposure to which can lead to different outcomes based on levels of vulnerability and resilience.

Remarkably, we found that these molecular markers that show distinct expression levels among vulnerable animals, additionally, exhibited strong correlations with the extent of behavioral vulnerability. We might have expected that any correlations with behavioral vulnerability would be negative for VLGUT and GAD metrics (because vulnerable animals present with an overall decrease in these markers), and positive for NPY. We did find a strong positive correlation for NPY, consistent with this expectation. However, contrary to expectation, we found that correlations of GAD and VGLUT metrics were also strongly positive among vulnerable animals, despite the group-wise decrease in vulnerable animals compared to resilient and sham controls. This suggests that while vulnerable animals present an overall decrease in GAD and VGLUT expression, the mechanisms associated with these pathways in individual vulnerable animals are more complex. Whether these molecular differences are preexisting in the individuals and lead to vulnerability, or emerge in vulnerable animals as a consequence of the injury, remains unclear.

### Predictors of vulnerability to anxiety dysfunction

We found, interestingly, that prior to injury, vulnerable animals spend less time in the open arms of the EZM, compared to the resilient and animals (EZM: F(2)=4.97, p=0.008), suggesting a more anxious phenotype. Following TBI, these vulnerable animals (but not the other groups) exhibited 2.5 to 3 fold increases in the proportion of time spent in the open arm, over their own baselines (Fig 1D). This anxiety profile post-injury in vulnerable animals remained robust, even after controlling for the baseline reduction.

Notably, the difference in EZM baselines is able to predict the post-injury behavioral outcome with approximately 70% accuracy (Fig 7C). Thus, behavioral measurements with the EZM are able to identify the subset of animals that will develop anxiety dysfunction upon being exposed to injury in the future. Understanding the neural basis of this difference in pre-injury behavior can give us clues to what makes an individual vulnerable. Specifically, examining molecular changes in the animals that show low EZM metrics at baseline will be a starting point to investigating pre-existing, causal molecular mechanisms predictive of vulnerability to anxiety following injury.

Our data found that about 13% of injured male C57 mice were vulnerable to anxiety dysfunction using a focal CCI model of injury. Other factors are also likely to influence the extent of vulnerability to anxiety dysfunction post-TBI, such as sex (Ashman et al., 2004, Kisser et al., 2017), genetic differences (Heilig et al., 1989, Cohen et al., 2008, Zhou et al., 2008), and environmental influences, including type of injury, disease, early life stress and chronic stressors (Vaishnavi et al., 2009, Bajwa et al., 2016, Nichols et al., 2016). The approach developed here sets the stage for exploring the effect of these other potential factors on behavioral vulnerability. In addition, it provides a general means to obtain a deeper understanding of neural mechanisms underlying vulnerability to anxiety dysfunction.

In this study, we have demonstrated that the effects of standardized injury are widely variable across individuals, and that vulnerability to anxiety dysfunction following TBI has specific neural signatures. These results establish that resilience and vulnerability are fundamental to understanding anxiety outcomes of TBI, and that accounting for individual variability can yield rich insights into neural mechanisms underlying post-injury anxiety outcomes.

## ACKNOWLEDGEMENTS

This work was supported by funding from the Johns Hopkins University (HA and SPM).

## AUTHOR CONTRIBUTIONS

HA and SPM designed the experiments, SPM and HA designed the analyses, JP performed the experiments and analyses, and HA, SPM and JP wrote the paper.

## METHODS

### Subjects

Adult male C57BL6J mice (Jackson Labs, Bar Harbor, ME, USA) were used in the experiments. Mice were housed in colonies of four (CCI and sham animals co-housed), in a 12 h light cycle (lights on from 7 am to 7 pm), with constant temperature and humidity (22°C and 40%). Food and water were available *ad libitum*. Animals’ weight was monitored weekly, averaging 32 g. Mice were 6–8 weeks-old at the beginning of experiments and allowed 2 weeks of acclimation before experiments began. Behavioral testing was conducted between 10 am and 5 pm. This study was carried out in accordance with the recommendations of IACUC guidelines. All experimental procedures were approved by Johns Hopkins Animal Care and Use Committee.

### Injury Procedures

Adult male mice underwent CCI (TBI group) or sham surgery (sham control group) at 6-8 weeks age. Animals were assigned to the two groups randomly. Experiments and analysis were performed in two independent batches (Cohort A: CCI 66, Sham 15; Cohort B: CCI 25, Sham 17; Total mice 103). Anesthesia protocols differed for the two cohorts: Cohort A: 3% isofluorene progressively reduced to 1%; Cohort B: Avertin (2,2,2 tribromoethanol - Sigma, St.Louis, MO) diluted in isotonic saline (500 mg/kg, i.p. After a midline skin incision, a circular craniotomy was made midway between Bregma and lambda with the medial edge of the craniotomy 0.5 mm lateral to the midline. The mice were then subjected to a moderate to severe CCI injury (Claus et al., 2010a, Wakade et al., 2010, Adwanikar et al., 2011, B. et al., 2017, Wang et al., 2018) using a convex impactor tip of 3 mm in diameter, using the following parameters: 4.5 m/s velocity, 1.50 mm depth of penetration and a sustained depression for 150 ms. After surgery, the scalp was closed with sutures, and mice were given 1 ml saline and 100 μls of 10% Meloxicam subcutaneously. Body temperature was maintained at 37°C with a warming blanket until full recovery from anesthesia (recovery of righting reflex). Sham-operated controls underwent the same surgical procedures with the exception of the traumatic injury. After recovery, mice were housed in individual cages for 72 h and monitored daily, then returned to their home-cages.

### Behavioral Tests and Apparatus

Baseline and post-injury behavioral testing were performed with all mice - TBI and sham - together, in the same order each week using three assays for testing anxiety-like behaviors. The Open-Field Test (OFT) consisted of a 40.6 cm × 40.6 cm sound-attenuating box. Mice were placed in the center of the field and allowed to freely explore for 20 min. The Elevated Plus Maze (EPM) consisted of two intersecting runways (50 cm × 50 cm × 5 cm), placed at 1 m from the ground. One runway had no walls (two open arms), while the other had 20 cm dark, high walls (two closed arms). Mice were placed in the center of the maze, facing one closed arm, and allowed to explore for 10 min. The Elevated Zero-Maze (EZM) consisted of a circular platform (width: 5 cm, inner diameter: 40 cm), placed at 1 m from the ground and divided in four quadrants. Two opposite quadrants had a 20 cm dark, high walls (closed arm), while the other two had no walls (open arms). Mice were placed facing the entrance of a closed arm, and allowed to explore for 10 min.

Testing proceeded as follows: animals were brought into the experimental room at least 30 min before the experiments began. They were first tested in the OFT, and given at least 2 h to recover before testing on the EZM. Twenty-four hours later, they were brought back to the experimental room and tested in the EPM. Mazes were cleaned with 70% ethanol between each animal. There were two trials before injury and four biweekly trials after injury. Baseline testing was performed 5-7 days apart, and injury was induced one week after the last baseline test. Baseline values were averaged for each animal to calculate their individual baseline level of anxiety.

Behavioral metrics for each trial were captured using an overhead camera and analyzed using Ethovision software (Noldus, Inc). Metrics analyzed include total distance traveled, distance traveled and time spent in defined zones (periphery vs center for OF; open vs closed arms for EPM and EZM), number of entrances in each arm (for EZM and EPM).

### Multidimensional behavior clustering clustering with validation (MBCV)

Behavioral metrics in the EZM and OFT on weeks one, three, five and seven, and EPM on weeks three, five and seven were combined into one 11-dimensional vector, referred to as the animals’ behavioral profile. (These metrics were normalized to their baselines.) PCA (Principal Components Analysis) was then applied to dataset consisting of the 11-dimensional behavioral profiles of all injured animals; this was done for each cohort independently (experiments in the cohorts were separated by ^~^1.5 years). The first four principal components (PCs) out of the 11 were chosen as the reduced-dimension representation of the dataset, because these four PCs accounted for the majority of the variability in the dataset (91% of the variability in Cohort A, and 87% in Cohort B). Following this dimensionality reduction, a clustering algorithm (k-means, with k=2) was applied *(kmeans* function in MATLAB) to identify two clusters of animals. The determination of whether animals in the two clusters exhibited distinct anxiety phenotypes was performed by the validation step in which the anxiety metrics from each of the three assays were compared between the two clusters. For the sham control animals, the above procedure was repeated on the 11-dimensional behavioral profiles of the sham animals from each cohort independently (just as for the TBI animals).

### Anatomical Metrics and Immunohistochemistry

Two weeks following behavioral testing, mice were deeply anesthetized (2.5% Avertin, 250 mg/kg body weight, i.p., Sigma, St. Louis, MO, USA) and transcardially perfused with 1 M phosphate buffer saline (PBS, 50 ml) followed by 4% paraformaldehyde (PFA, 100 ml, Sigma, St. Louis, MO, USA). Brains were removed and post-fixed in 4% PFA (50 ml) for 72 h, then transferred to a solution of 30% sucrose in 4% PFA, where they were kept refrigerated until sectioning. Coronal sections (40 μm) of the whole brain were made using a slide microtome (Leica Microsystems, model CM-1860).

The primary antibodies used and their concentrations were: anti-GAD65/67 (1:1000, EMD Millipore), anti-VGLUT (1:1000, Thermofisher), and anti-Neuropeptide-Y (1:000, ABCAM). Mounted sections were washed 3 times for 10 min each wash in 0.1M PBS and then incubated in blocking solution. For GAD65/67 and VGLUT, we used a blocking solution containing 10% normal goat serum and 0.5% Triton X-100 in PBS for 2 h at room temperature. For NPY, sections were blocked with a Mouse on Mouse immunodetection kit (Vector Labs) to reduce endogenous staining, for 1h at room temperature. Slides were then incubated with primary antibody, in 10% normal goat serum for GAD65/67 and VGLUT, and 5% mouse IgG blocking reagent for NPY, in 0.5% Triton X-100, for 72 h at 4° C. After incubation, the sections were rinsed three times in PBS for 10min, and incubated with the secondary antibody, with the appropriate blocking serum, 0.5% Triton X-100, for 2 h in room temperature. The secondary antibodies used were Alexa Fluor 488 (1:1000, Abcam) for GAD6567 and VGLUT1, and Alexa Fluor 405 (1:1000, Abcam) for NPY. Sections were washed three times for 10 min with PBS, and incubated for 30 min with NeuroTrace 640/660 Deep-Red Fluorescent Nissl (Thermofisher) and mounted with Vectashield antifade mounting medium (Vector Labs). Separated, consecutive sections were used for each antibody. Fluorescent images of the BLA, mPFC and vHPC were taken with a Zeiss LSM 700 microscope (Carl Zeiss, Germany) using 40x objectives. Four bilateral immunofluorescent labeled sections were taken in each mouse (8 images per animal per region, per marker), sections were approximately 200 um apart. Images were pre-processed using FIJI, and then the number and intensity of puncta were calculated using the IMFLAN3D analysis package (Tai et al., 2007, Schindelin et al., 2012, Ha et al., 2018). (A ‘punctum’ was defined as a cluster of ‘connected’ pixels identified in an objective, automated manner using the *bwlabeln* function in MATLAB with the ‘eight-connected neighborhood’ criterion (Tai et al., 2007, Schindelin et al., 2012, Ha et al., 2018))

For volumetric measurements, fluorescent images of the brain in the peri-injury region were taken with an Axio-zoom v16 microscope (Carl Zeiss, Germany) using a 10× objective. Volume estimation was performed as described previously (Claus et al., 2010b), adopting Cavalieri estimation (Gundersen and Jensen, 1987). Images from four sections were taken for each mouse in each area. We determined the position of each section by using Mouse Brain Atlas coordinates (Paxinos and Franklin, 2004). Sections were located between −0.70 and −2.46 mm Bregma, and they contained the lesioned area in the injured mice or equivalent position in sham controls. BLA images were taken between −1.22 and 2.18 mm Bregma. Images were pre-processed using Fiji software (Schindelin et al., 2012). The hemispheric and BLA areas were outlined using Fiji on the ipsilateral and contralateral sides of the four brain sections, volume was calculated as the sum of the areas multiplied by the distance between sections (300 μm). The ipsilateral volume was normalized to the contralateral volume to estimate the extent of hemispheric volume loss.

### Analyses on baseline measurements

For the ‘baseline equalized analysis (Fig. 7B), the average value of the EZM anxiety metrics measured prior to injury for the resilient and sham groups were artificially adjusted to match the average value of this metric for vulnerable animals. Following this ‘equalization of means, the post-injury outcomes were normalized to these adjusted baseline values and were compared among the three groups.

A standard receiver operating characteristic, or ROC, analysis was performed on the baseline values of the EZM anxiety metric to examine how well these baseline values could predict vulnerability to subsequent injury (‘true positives’; Fig. 7C). The area under the ROC curve (AUC) provided an estimate of the predictive power.

### Statistical analyses

For statistical analysis of the behavioral results, group means were compared using ANOVA (2-way or 3way, unbalanced, repeated measures). Post-hoc testing was performed with paired t-tests between groups followed by the Holm-Bonferroni multiple comparisons test, abbreviated as ‘HBMC test’ in the text.

For statistical analysis of immunohistochemical results, group means were compared using three-way ANOVA. Post-hoc testing was performed with paired t-tests between groups followed by HBMC test. For comparisons that showed significant post-hoc differences, we calculated effect size (*η^2^ or Eta-squared*) to quantify the magnitude of the effect. It was calculated using the Effect Size toolbox v1.6.1 (Hentschke and Stüttgen, 2011), and the function *mes2way,* in MATLAB, following the formula: η^2^=SS_effect_/SS_total_. We report the strength of effect sizes following standard convention as: small (η^2^ = 0.02), medium (η^2^ = 0.06), and large (η^2^ = 0.14) (Cohen, 2013).

In all the instances where the metric showed that the vulnerable animals were significantly different from resilient or sham controls, the correlation between the molecular metric and behavioral metric of anxiety (EZM, week seven or week 5) was computed as Pearson’s correlation, using the *corr function in* MATLAB. We adopt the criteria of r>0.6 for a strong correlation, and 0.2<r<0.6 for a weak to moderate correlation (Lakens, 2013).

For statistical analysis of volumetric measurements, group means were compared using ANOVA, and post-hoc t-tests followed by HBMC test.

