## Supplementary figures and tables for "Neural markers of vulnerability to anxiety outcomes following traumatic brain injury"

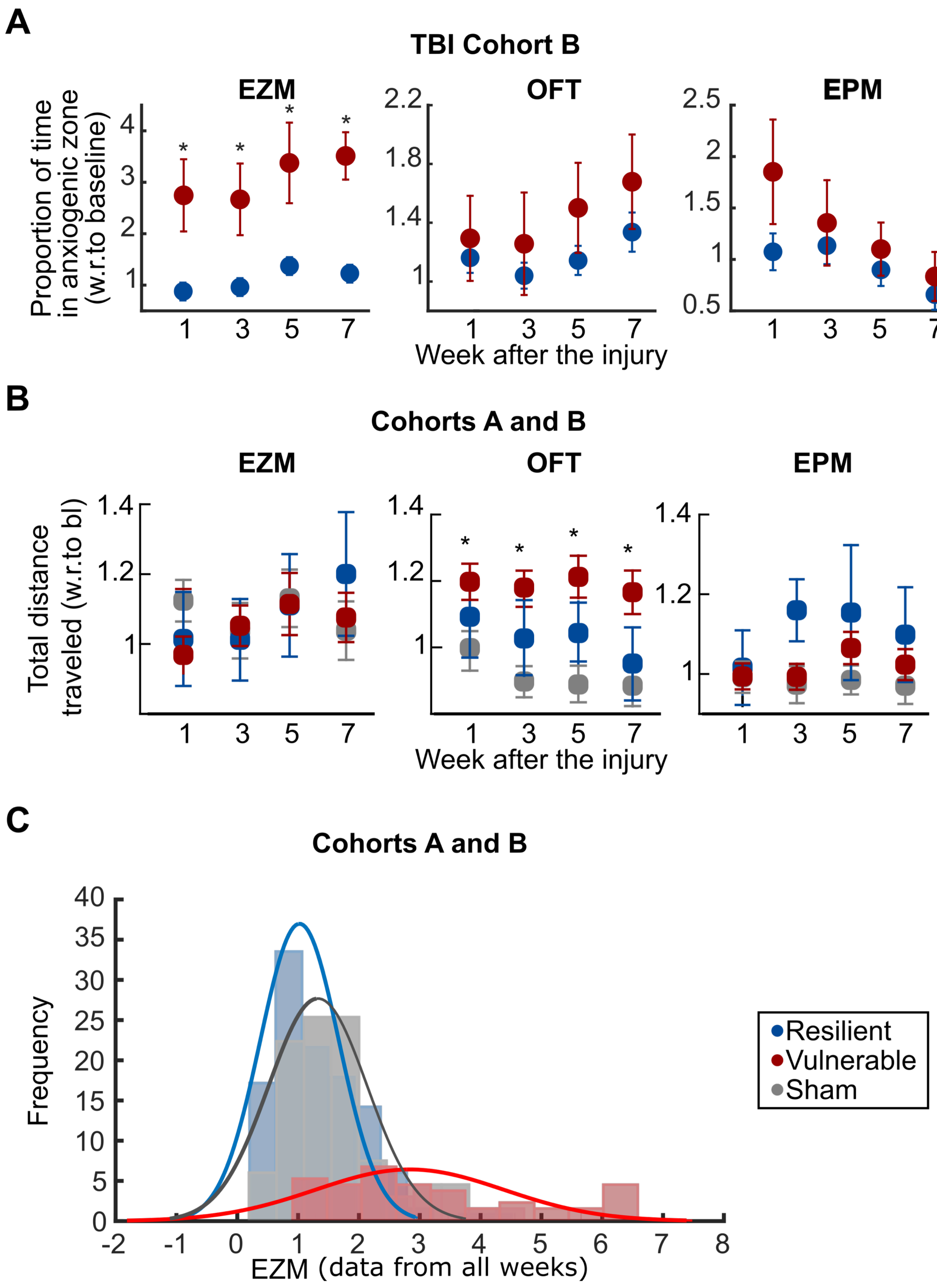


**Supplementary Figure 1. TBI animals present distinct behavioral profiles of vulnerability versus resilience to anxiety following TBI.** ***(Related to Figure 2.)***

**(A)** Validation step after multidimensional behavioral clustering (Cohort B). Plots of anxiety metric from each assay, comparing the red versus blue clusters of TBI animals from Cohort B. ***Left***: EZM; red animals show decreased anxiety (greater proportion of time in anxiogenic zone) after injury compared to blue animals (2-way ANOVA: F=56.34, p=<0.001, main effect); '*': p<0.05, posthoc testing by week of red vs. blue followed by HBMC test. Results show that red animals in Cohort B exhibit similar behavioral phenotype to red animals in Cohort A. ***Middle***: OFT; 2-way ANOVA: F=2.08, p=0.1, marginal main effect. ***Right***: EPM; 2-way ANOVA: F=1.43, p=0.24, no main effect. All three panels: Shown are mean +/- s.e.m.
**(B)** Total distance travelled by animals in each of the assays; Cohorts A and B combined. ***Left***: EZM; no main effect (3-way ANOVA, F=xx, p=xx). ***Middle***: OFT; vulnerable animals traveled more than sham animals 3-way ANOVA, F=29.74, p=0, main effect; '*': p<0.05, posthoc testing by week of vulnerable animals vs. sham controls followed by HBMC test. ***Right***: EPM; 3-way ANOVA: F=95, p=0, main effect of group; no post-hoc effects.
**(C)** Histograms showing the proportion of time spent in the open arms of the EZM; datapoints from all four time-points combined. Histograms shown separately for vulnerable (red), resilient (blue) and sham (grey) animals. Curves represent best-fitting Gaussians (using the *nlinfit* function in MATLAB). Results show substantial (nearly total) overlap between resilient and sham animals, supporting the hypothesis that comparing behavioral outcomes in the sham group versus the TBI group (vulnerable + resilient) can mask effects of TBI on vulnerable animals and confound interpretation (see text).

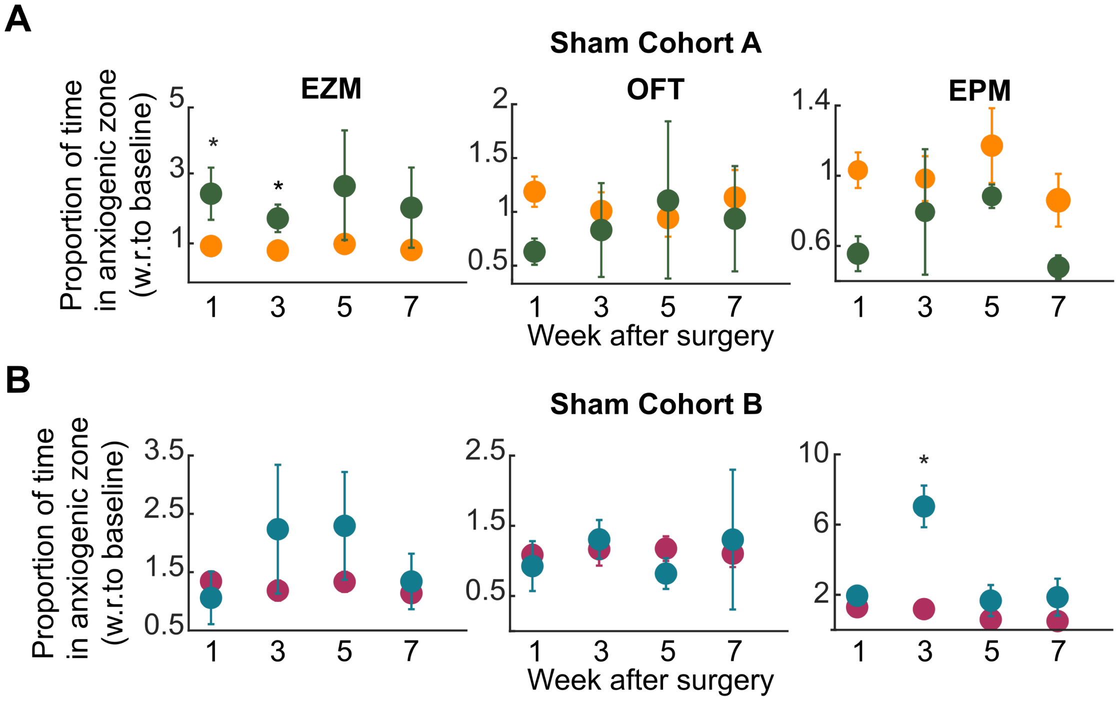


**Supplementary Figure 2. Clustering sham animals does not produce behaviorally distinct groups.** ***(Related to Figure 3.)***
**(A,B)** Validation step after multidimensional behavioral clustering applied to behavioral profiles of sham animals from Cohort A (A) and Cohort B (B). Plots of anxiety metric from each assay, comparing the green versus orange clusters of sham animals from Cohort A (A), and teal versus maroon clusters of sham animals from Cohort B (B). ***Left***: EZM; there was a main effect of cluster for Cohort A (A, 2-way ANOVA, F=40.48, p<0.0001) and cohort B (B, 2-way ANOVA, F=4.36, p=0.004), with a post-hoc effect on weeks one

and three for cohort A (‘*’: green cluster > orange cluster; posthoc t-tests followed by HBMC test). ***Middle***: OFT; No main effect of cluster for either cohort A (2-way ANOVA, F=0.58, p=0.44) or cohort B (2-way ANOVA, F=0.003, p=0.85). ***Right***: No main effect of cluster for cohort A (2-way ANOVA, F=2.87, p=0.09), but there was a main effect of cluster for cohort B (2-way ANOVA, F=60.63 p<0.001), with a post-hoc effect on week three (‘*’: teal cluster > maroon cluster; posthoc t-tests followed by HBMC test).

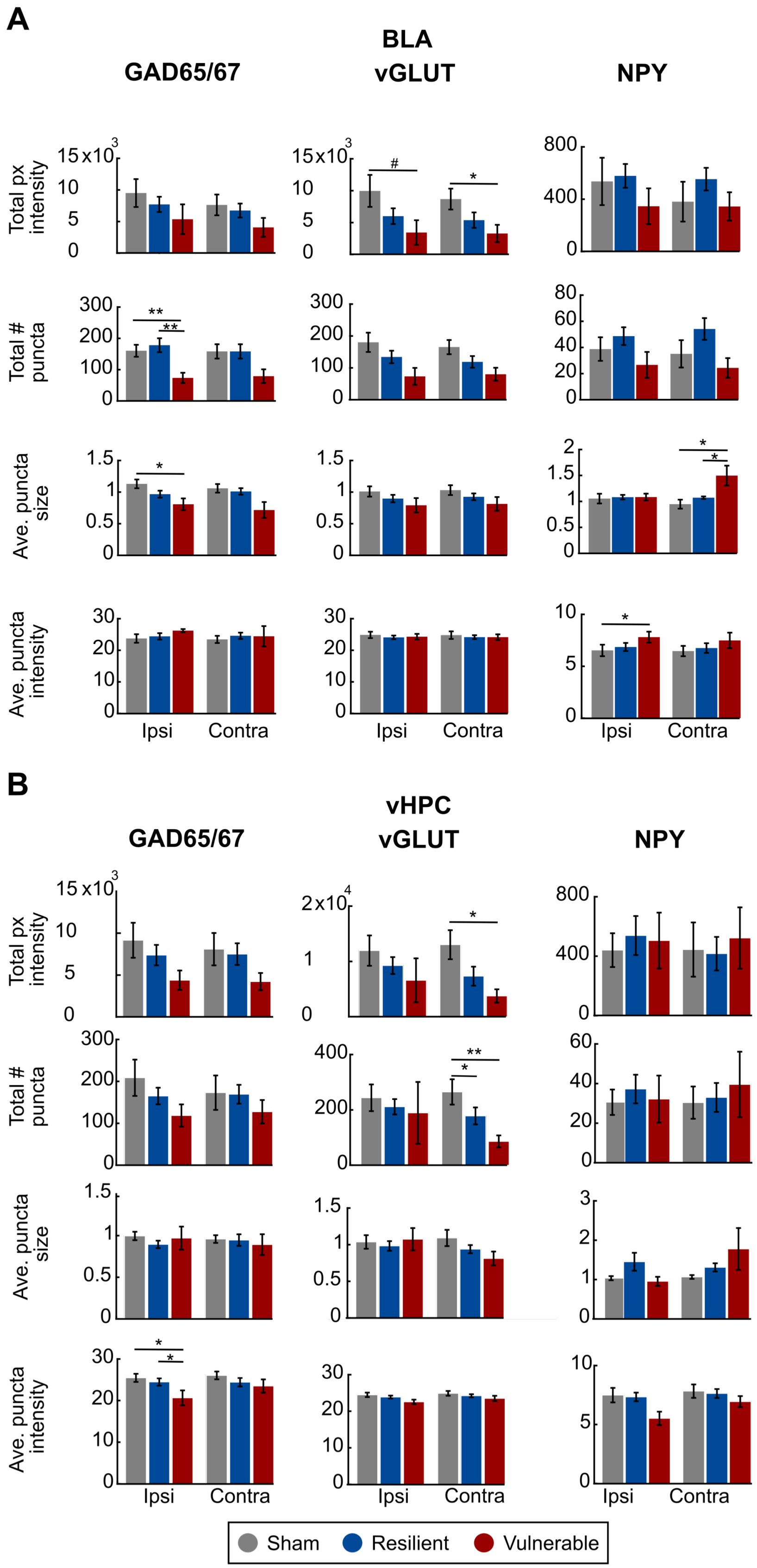

A
BLA
GAD65/67
vGLUT
NPY
Total px intensity
Total # puncta
Ave. puncta size
Ave. puncta intensity
Ipsi
Contra
B
vHPC
GAD65/67
vGLUT
NPY
Sham
Resilient
Vulnerable

**Supplementary Figure 3. Vulnerable animals present significant differences in molecular markers compared to resilient and sham animals in the BLA and vHPC.** ***(Related to Figure 5).***

**(A,B)** Comparison of various metrics of immunostaining (rows) for each of the three molecular markers (columns) in BLA (A) and vHPC (B). Shown are mean +/- s.e.m; data from 53 animals (vulnerable= all 9, resilient=28, sham=16; resilient and sham animals for immunostaining were selected randomly from their respective groups) *: $p<0.05$ and **: $p<0.01$, for difference between groups: ANOVA followed by post-hoc t-tests with HBMC test.

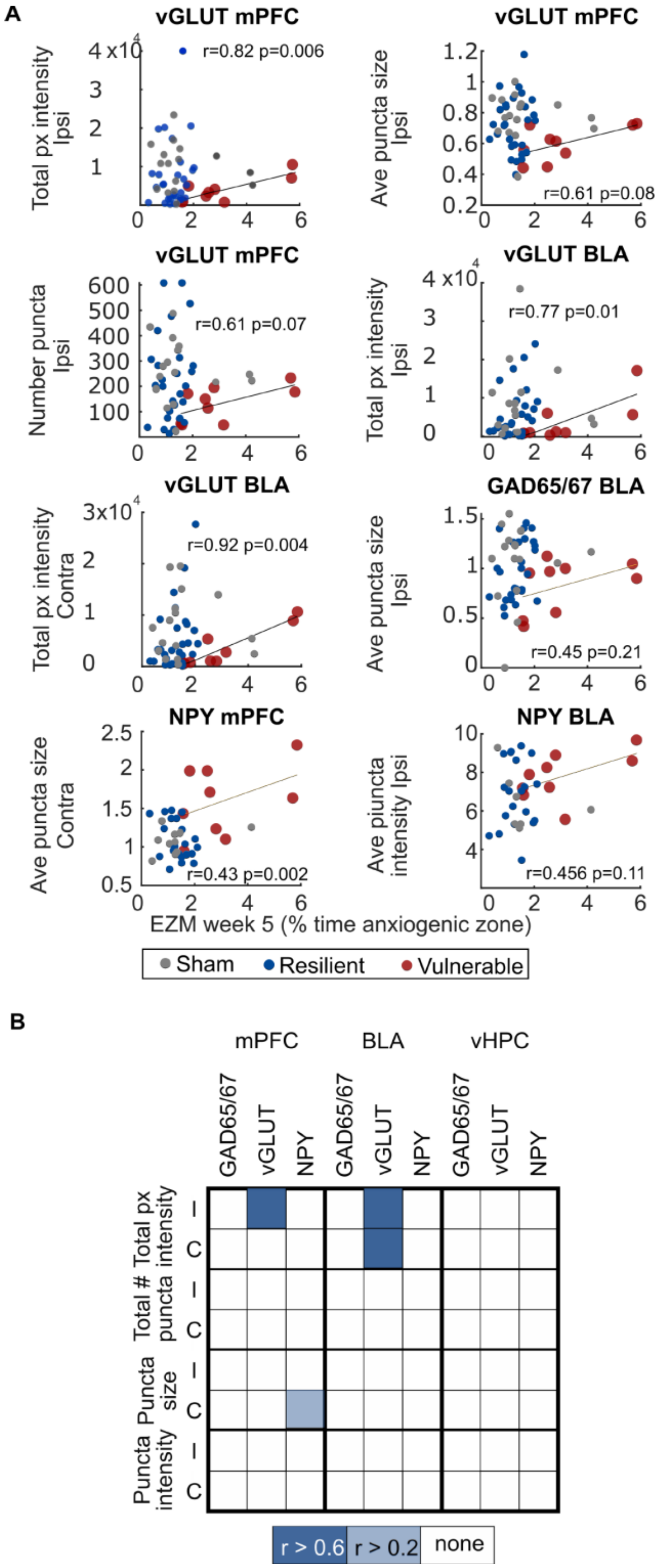
A
vGLUT mPFC
Total px intensity Ipsi
r=0.82 p=0.006
vGLUT mPFC
Ave puncta size Ipsi
r=0.61 p=0.08
vGLUT mPFC
Number puncta Ipsi
r=0.61 p=0.07
vGLUT BLA
Total px intensity Ipsi
r=0.77 p=0.01
vGLUT BLA
Total px intensity Contra
r=0.92 p=0.004
GAD65/67 BLA
Ave puncta size Ipsi
r=0.45 p=0.21
NPY mPFC
Ave puncta size Contra
r=0.43 p=0.002
NPY BLA
Ave piuncta intensity Ipsi
r=0.456 p=0.11
EZM week 5 (% time anxiogenic zone)
Sham
Resilient
Vulnerable
B
mPFC
BLA
vHPC
GAD65/67
vGLUT
NPY
Total px intensity
Total # puncta
Puncta size
Puncta intensity
I
C
r > 0.6
r > 0.2
none

**Supplementary Figure 4. Correlation between molecular metrics and anxiety outcome among vulnerable animals generalizes to another anxiety metric (EZM, week 5)** ***(Related to Figure 6).***
This figure tests for generality of the conclusion that molecular metrics correlate with a behavioral metric of anxiety (EZM, week 7) specifically in vulnerable animals. In order to identify an alternative behavioral variable in a principled manner, we reasoned that the behavioral variable that contributed most to the first principal component would be a sensible one. To this end, we ranked the original 11 variables by their similarity (loading) with the first principal component (PC1), and found that the proportion of time spent in the open arms of EZM in week 5 had the largest similarity with PC1 (loading: Cohort A: 0.13, Cohort B: 0.52, %loading relative to those of other variables: Cohort A: 21.4%, Cohort B: 22.9%). We then performed correlation analyses between the molecular metrics in vulnerable animals and the metric of anxiety measured in EZM on week 5. Results show strong positive correlations specifically for vulnerable animals (but not resilient or sham animals), similar to Figure 6, thereby supporting the generality of the result.
**(A)** Scatter plots of molecular metrics versus anxiety metric (proportion of time spent in the anxiogenic zone) measured on week 5 in the EZM. Each dot denotes one mouse; red: vulnerable, n= all 9 animals; blue: resilient, n=28 (GAD65/67), n=29 (vGLUT), n=23 (NPY); grey: sham, n=16 (GAD65/67), n=17 (vGLUT), n=12 (NPY); r: Pearson's correlation, correlation, line: best-fit line, p: p-value from Pearson's correlation test. All other conventions as in Figure 6A.
**(B)** Visualization of the strength of correlations in A. The matrix represents the strength of correlations between the proportion of time spent by vulnerable animals in the anxiogenic zone of EZM in week 5, against each of various molecular metrics (rows) of GAD65/67, VGLUT and NPY (columns) measured in the three brain regions in the ipsi (I) and contra (C) hemispheres. All correlation values indicated are positive.

| | Mean contralateral volume (e+07) | SEM contralateral volume (e+06) | Mean ipsilateral volume (e+07) | SEM ipsilateral volume (e+06) |
|---|---|---|---|---|
| **Sham controls** | 2.3 | 2.8 | 2.2995 | 2.9436 |
| **Resilient** | 1.65 | 2.44 | 1.2873 | 2.1918 |
| **Vulnerable** | 1.64 | 5.14 | 1.2708 | 4.0155 |

**Supplementary Table 1. Contralateral and ipsilateral hemispheric volume in sham controls, resilient and vulnerable animals.** ***(Related to Figure 2).***
A three-way ANOVA shows no main effect of treatment in the contralateral side ($F=1.52$, $p=0.23$), and a main effect of treatment in the ipsilateral side ($F=4.42$, $p=0.019$), and TBI animals (resilient and vulnerable) presented a significant decrease in ipsilateral hemispheric volume compared to sham controls ($p<0.05$). Resilient and vulnerable animals did not differ from each other.
